# MR-Clust: Clustering of genetic variants in Mendelian randomization with similar causal estimates

**DOI:** 10.1101/2019.12.18.881326

**Authors:** Christopher N Foley, Paul D W Kirk, Stephen Burgess

**Affiliations:** University of Cambridge; MRC Biostatistics Unit

## Abstract

**Motivation:** Mendelian randomization is an epidemiological technique that uses genetic variants as instrumental variables to estimate the causal effect of a risk factor on an outcome. We consider a scenario in which causal estimates based on each variant in turn differ more strongly than expected by chance alone, but the variants can be divided into distinct clusters, such that all variants in the cluster have similar causal estimates. This scenario is likely to occur when there are several distinct causal mechanisms by which a risk factor influences an outcome with different magnitudes of causal effect. We have developed an algorithm MR-Clust that finds such clusters of variants, and so can identify variants that reflect distinct causal mechanisms. Two features of our clustering algorithm are that it accounts for uncertainty in the causal estimates, and it includes ‘null’ and ‘junk’ clusters, to provide protection against the detection of spurious clusters.

**Results:** Our algorithm correctly detected the number of clusters in a simulation analysis, outperforming the popular Mclust method. In an applied example considering the effect of blood pressure on coronary artery disease risk, the method detected four clusters of genetic variants. A hypothesis-free search suggested that variants in the cluster with a negative effect of blood pressure on coronary artery disease risk were more strongly related to trunk fat percentage and other adiposity measures than variants not in this cluster.

**Availability and Implementation:** MR-Clust can be downloaded from https://github.com/cnfoley/mrclust.

**Supplementary Information:** Supplementary Material is included in the submission.

## 1 Introduction

Genome-wide association studies have discovered many genetic variants associated with various traits and conditions. Such genetic variants can aid understanding of the biological mechanisms that influence traits [1]. They can also be used to link modifiable traits to disease outcomes. Due to random mating (that is, choice of partner is independent of the genetic variants under investigation) and Mendel’s laws of segregation and independent assortment, genetic variants are typically distributed independently of traits that they do not directly influence, and so can be treated similarly to random treatment assignment in a randomized controlled trial [2, 3]. Genetic variants associated with a given trait are therefore plausible instrumental variables (IVs) for that trait [4]. The use of genetic variants as IVs to assess the causal effect of a risk factor on an outcome is known as Mendelian randomization [5].

While the hypothesis of whether a risk factor has a causal effect on an outcome can be assessed with a single valid IV [4], most genetic variants do not explain enough variability in the risk factor to have sufficient power to reliably detect a moderate-sized causal effect. Additionally, it is prudent to use all relevant data to address the causal hypothesis of interest. Under strict parametric assumptions (described below), the causal estimates based on each valid IV will target the same causal parameter – the average causal effect [6]. Excess heterogeneity between causal estimates from different genetic variants is often interpreted as evidence that not all genetic variants are valid instrumental variables [7].

However, it may be that different genetic variants influence the risk factor in distinct ways, leading to heterogeneity between causal estimates calculated using different variants. For example, several hundred genetic variants have been demonstrated to be independently associated with blood pressure [8]. Different genetic variants may influence blood pressure via distinct biological mechanisms. Alternatively, some variants may influence traits that are causally upstream of blood pressure rather than blood pressure directly. Or it may be that blood pressure is in fact a composite trait consisting of multiple components that is captured only as a single measurement. Variants that influence the risk factor in a similar way are likely to have similar causal estimates.

Several previous attempts have been made to cluster genetic variants that are associated with a given risk factor. Walter et al. [9] took 32 genetic variants associated with body mass index (BMI) and divided the variants into four groups based on biological understanding of the function of the variants. They then compared the causal estimates of BMI on depression based on each group of variants. Udler et al. [10] took 94 variants associated with Type 2 diabetes, and divided the variants into 7 groups based on the their associations with 47 diabetes-related traits. Tanigawa et al. [11] applied a truncated singular value decomposition method to genetic association estimates from the UK Biobank study to find clusters of variants having similar associations with a range of traits.

In this paper, we introduce a method to cluster variants that have similar causal estimates for the given risk factor and outcome. As we do not use data on genetic associations with alternative traits to form the clusters, an advantage of this approach is that genetic associations with traits can be used to validate the division into clusters. If traits can be found that predict cluster membership, this increases the plausibility that the clusters have a biological interpretation. We refer to our method as MR-Clust.

Our manuscript is structured as follows. First, we provide an overview of Mendelian randomization, and introduce the modelling assumptions and notation used in the manuscript (Section 2). We also consider factors that may lead to heterogeneity between causal estimates based on different genetic variants, and in particular investigate how this would lead to clustered heterogeneity. Next, we introduce a statistical approach for detecting clusters of variants with similar causal estimates, which are likely to influence the risk factor in a similar way (Section 3). There are two distinct aspects of our method over conventional applications of clustering. First, we account for uncertainty in the causal estimates that we are clustering. Secondly, we include a ‘junk’ cluster in our model, so that variants with estimates that do not fit into any clusters are included in the junk cluster rather than any other cluster. We apply our method in a simulation study, and to consider 180 independent genetic variants associated with blood pressure at a genome-wide level of significance, and find clusters in the causal estimates of blood pressure traits on coronary artery disease (CAD) risk (Section 4). We conclude by discussing the results of the manuscript, and their application to epidemiological practice (Section 5).

## 2 System and methods

The aim of a Mendelian randomization analysis is to establish whether there exists a causal relationship between a risk factor *X* and an outcome *Y* using genetic variants *G*_*j*_, *j* = 1, 2, …, *J* as instrumental variables. An additional aim is to estimate the causal effect of the risk factor on the outcome. In this section, we introduce assumptions and methods for IV estimation, and discuss when the estimates based on different IVs will be similar and when they will be different.

### 2.1 Instrumental variable assumptions

A genetic variant *G*_*j*_ is a valid instrumental variable if it satisfies three assumptions:

- (relevance) it is associated with the risk factor,
- (exchangeability) its association with the outcome is not confounded, and
- (exclusion restriction) it has no effect on the outcome except that mediated via the risk factor [12, 13].

Under these assumptions, any association between the genetic variant and the outcome is indicative of a causal effect of the risk factor on the outcome [14].

To estimate a causal parameter, we make further parametric assumptions of linearity and homogeneity in the relationships between the genetic variant, risk factor, and outcome. Specifically:

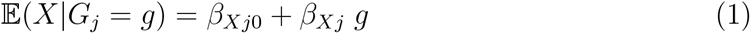

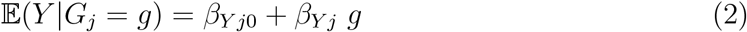

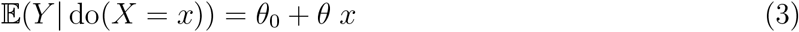

where *θ* is the average causal effect of the risk factor on the outcome [15], and do(*X* = *x*) is Pearl’s do operator, meaning that the risk factor is intervened on to take value *x* [16]. This model can be illustrated as a directed acyclic graph (Figure 1).

**Figure 1:**
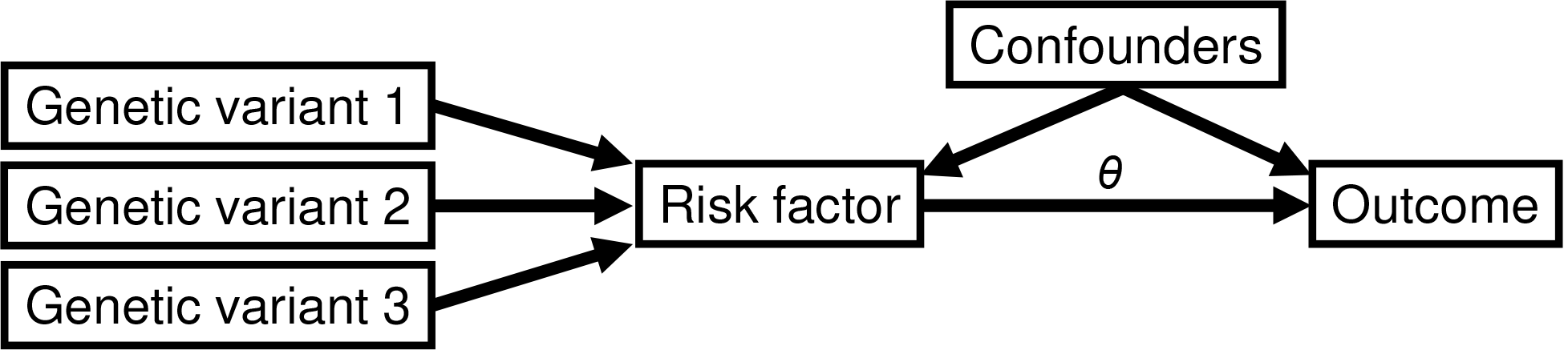
Directed acyclic graph illustrating relationships between three genetic variants that are valid instrumental variables with a risk factor, outcome, and confounders of the risk factor–outcome associations. The causal effect of the risk factor on the outcome is indicated by *θ*.

It can be shown that *θ* can be estimated consistently as the ratio of the estimated genetic association with the outcome divided by the estimated genetic association with the risk factor [17]:

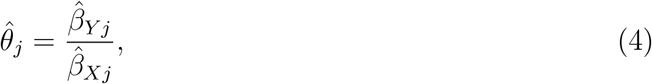

which we call the ratio estimate of the *j*th variant. The standard error of this quantity 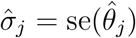 can be estimated using the delta method:

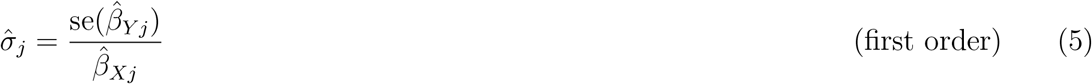

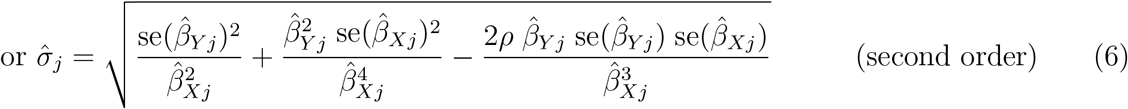

where *ρ* is the correlation between the genetic association estimates 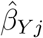 and 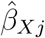.

We note that these parametric assumptions are sufficient, but not necessary for the estimation of the average causal effect; weaker assumptions have been proposed [18]. Alternatively, under the monotonicity assumption (the genetic variant increases the risk factor in all individuals in the population, or decreases the risk factor in all individuals), a local average causal effect can be estimated [19]. However, local average causal effects may differ between valid IVs. We return to this point in the discussion.

### 2.2 Heterogeneity between causal estimates and clustered heterogeneity

Even if the IV assumptions and parametric assumptions (1) and (2) are satisfied for each genetic variant, it is plausible that the variant-specific ratio estimates 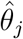 differ by more than expected due to chance alone. We are particularly interested in the case where there are distinct values of the causal effect that are evidenced by multiple genetic variants, such that if the sample size were to tend towards infinity, the ratio estimates would tend towards a number of distinct values. We refer to this situation as clustered heterogeneity. Clustered heterogeneity is interesting to investigate as the identity of the genetic variants in the clusters may reveal information about the risk factor and how it relates to the outcome. Figure 2 illustrates how different variants may be associated with the risk factor and outcome via different mechanisms. This situation could arise in a number of ways:

1. Risk factor is a composite trait: The risk factor is not a single entity, but in fact contains multiple components with distinct causal effects. For example, although serum cholesterol concentration can be expressed as a single measurement, evidence suggests that cholesterol carried by low-density lipoprotein particles has a different causal relationship to CAD risk compared with cholesterol carried by high-density lipoprotein particles [20].
2. Multiple versions of treatment: The risk factor can be intervened on in different ways, and each intervention leads to a different size of change in the outcome. For example, interventions to lower BMI via decreasing an individual’s caloric intake are likely to lead to less cardiovascular benefit compared with interventions to increase metabolic rate.
3. Multiple biological pathways: Even if the risk factor is a single trait and there is a single version of treatment, genetic variants may associate with the risk factor via different biological pathways, which may influence the outcome directly (that is, not via the risk factor).

**Figure 2:**
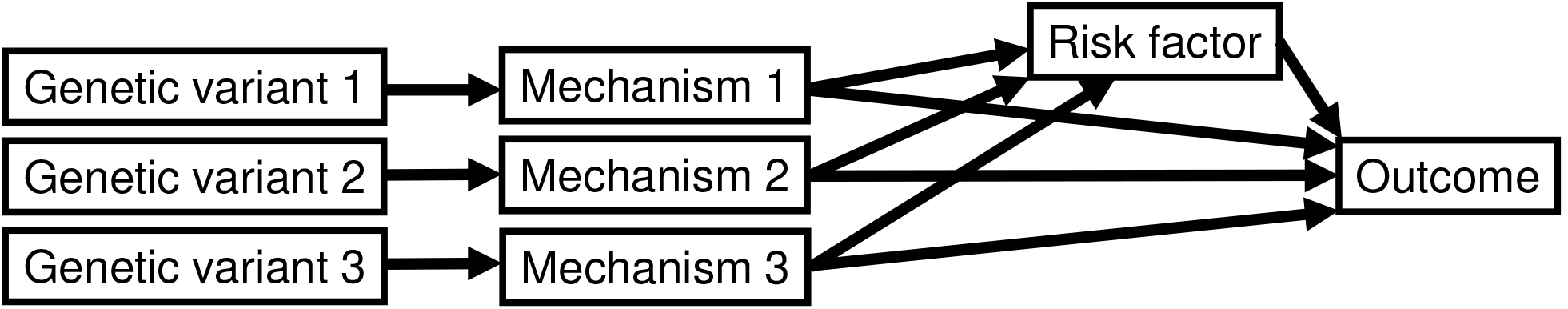
Scenario that could lead to clustered heterogeneity, defined as the case where causal estimates from multiple variants tend towards a number of distinct values as the sample size increases. Clustered heterogeneity could arise in a number of ways: the mechanisms may represent distinct components of the risk factor, or distinct pathways by which the risk factor may influence the outcome, or intermediaries on the causal pathway from the genetic variant to the outcome.

In the first two situations, identifying features of genetic variants in different clusters could help explain how the outcome is influenced by different components of the risk factor or different causal pathways from the risk factor, and hence inform our biological understanding of the causal relationship between the risk factor and outcome. In the third situation, the IV assumptions are violated, as the effects of the genetic variants on the outcome are not completely mediated via the risk factor. In this case, investigating traits that associate preferentially with variants in different clusters could identify intermediaries on the relevant causal pathway for each cluster.

In Appendix Section A, we provide some theoretical motivation that clustered heterogeneity arises if and only if genetic variants in the same cluster affect the outcome via the same distinct causal pathway, under assumptions of linearity and homogeneity.

## 3 Algorithm

We proceed to introduce a statistical method for clustering causal estimates from different genetic variants. We suppose that there are *K* + 2 disjoint clusters of genetic variants: *K* substantive clusters, a null cluster, and a junk cluster. The substantive clusters *S*_1_, …, *S*_*K*_ have means *θ*_*k*_, *k* = 1, …, *K*. The null cluster *S*_0_ has mean *θ*_0_ = 0. The presence of the null cluster ensures that genetic variants which do not suggest a causal effect of the risk factor do not contribute to the estimates of the substantive cluster means. The junk cluster *S*_*K* +1_ comprises all remaining genetic variables that are not members of the other clusters. The presence of the junk cluster ensures that genetic variants which do not fit into any of the substantive clusters do not contribute to the estimates of the substantive cluster means. Together, the null and junk clusters require there to be substantial evidence of similarity of estimates from several genetic variants to define a substantive cluster. This should minimize false positive findings from the method.

### 3.1 Mixture model

For each genetic variant *j* = 1, …, *J*, we introduce a cluster allocation label *z*_*j*_, such that *z*_*j*_ = *k* ⟺ *G*_*j*_ ∈ *S*_*k*_. For variants in the substantive and null clusters, we assume that the ratio estimate 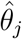 for variant *j* in cluster *k* follows a normal distribution with mean *θ*_*k*_ and standard deviation 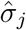, taken as the standard error of the *j*th ratio estimate:

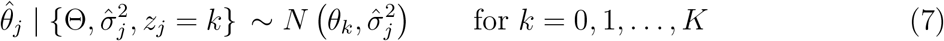

where Θ is a vector of the cluster means. For simplicity, we do not account for uncertainty in the estimate of the standard error.

Following Crook et al. [21], we assume ratio estimates for variants in the junk cluster follow a generalized t-distribution with degrees of freedom *ν* = 4, mean *μ* taken as the sample mean of all the ratio estimates 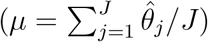, and scale parameter *ψ* taken as

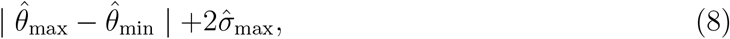

where 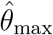 is the maximum of the ratio estimates, 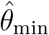 is the minimum of the ratio estimates, and 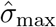 is the maximum of the standard errors of the ratio estimates. We discuss the specification of this distribution in Appendix Section B.

We obtain a mixture model for the ratio estimates 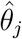:

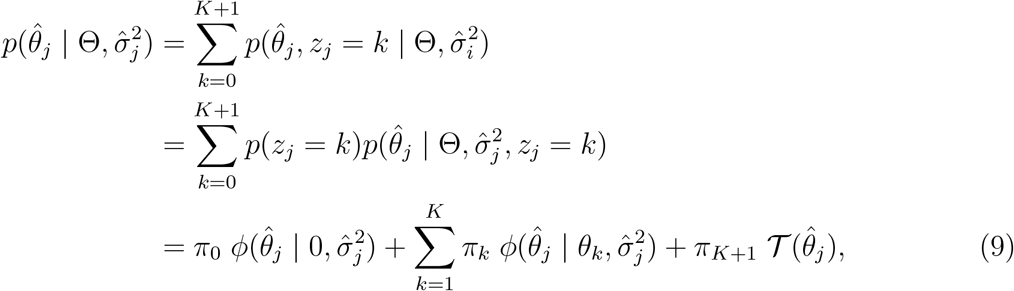

where *π*_*k*_ is the mixture proportion for cluster *k*, *φ*(*x* | *μ, σ*^2^) denotes the univariate normal density evaluated at *x* with mean *μ* and variance *σ*^2^, and 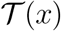 denotes the generalized t-distribution evaluated at *x* with degrees of freedom *ν* = 4, and mean *μ* and scale parameter *ψ* as discussed above.

### 3.2 Parameter estimation via expectation maximization

The 2*K* + 1 parameters *θ*_*k*_ and *π*_*k*_ (*K* cluster means and *K* + 2 proportions, less one as the proportions must sum to one: 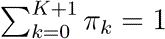) in equation (9) are estimated via an expectation-maximization (EM) algorithm for a given number of substantive clusters *K*. We then estimate the number of substantive clusters.

The log-likelihood of the sample data 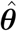 (the ratio estimates) is

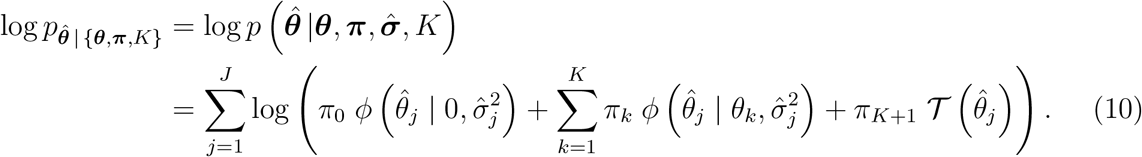

We denote the maximum likelihood estimate (MLE) of the unknown parameters for a given *K* as 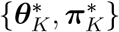. For ease of presentation, we drop the index *K* in this section.

We index each iteration of the EM algorithm by the variable *i* so that the pair {***θ***^(*i*)^, ***π***^(*i*)^} denotes estimates of the cluster means and mixture proportions at the *i*th iteration of the algorithm. We stop updating the parameters when the difference in log-likelihood between two iterations falls below a user-defined tolerance *δ*.

We describe the algorithm in three main steps: (i) an initialization step to obtain initial values of the parameters, and (ii) an expectation step and (iii) a maximization step to update the parameter values.

#### Initialization step

Reliable estimation of the MLE might depend crucially on the initialization of the parameters. To mitigate sensitivity to the initialization, our algorithm computes multiple estimates of the MLE over various initializations of the parameters. When *K* > 0, for each initialization we generate values for the cluster means {****θ****^(0)^} via a *k*-means clustering of the data 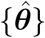. We note this method does not account for the uncertainty in the ratio estimates. The initial mixture proportions {****π****^(0)^} are computed by first randomly drawing values for the proportion of samples in the null and junk mixtures 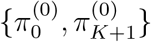 from the range (0.05, 0.4). This ensures that the prior probability of belonging to either the null or junk cluster is at least 10% and at most 80%. The remaining parameters are 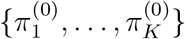 then computed as the proportion of observations a)signed to each of the *K* clusters from the *k*-means analysis multiplied by 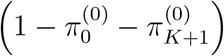.

#### Expectation step

Let ****Z**** denote the collection of cluster allocation labels {*z*_1_, *z*_2_, …, *z*_*J*_} for the variants *j* = 1, 2, …, *J*. Before updating the unknown parameters in the maximization step, we first evaluate:

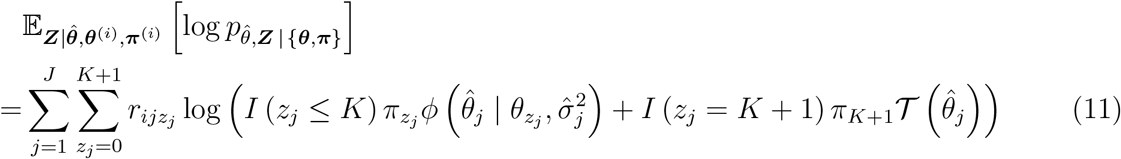

which requires computation of the conditional allocation probabilities for each observation:

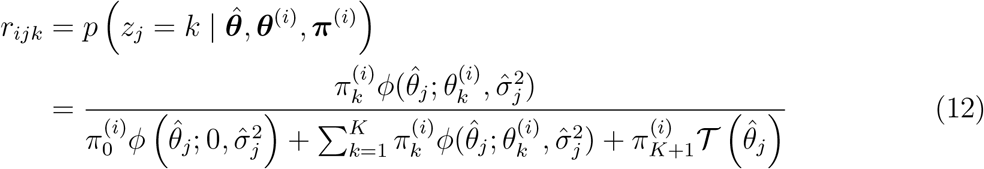

for *k* = 1, …, *K*, and:

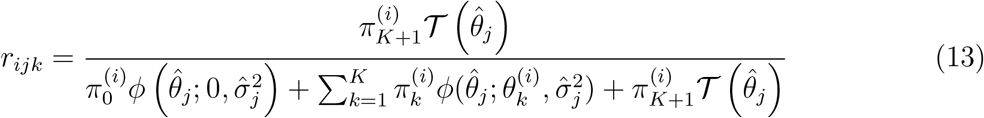

for *k* = *K* + 1. The *r*_*ijk*_ are sometimes referred to as the responsibilities of the *k*th component for the *j*th observation (evaluated here at the *i*th iteration of the EM algorithm).

#### Maximization step

Updates for the unknown parameters are obtained by maximizing equation (11). For the cluster means *θ*_*k*_, we solve the system of equations

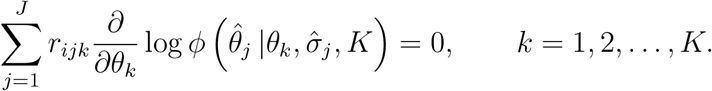

Re-arranging for *θ*_*k*_, and taking this as the update 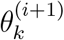, returns

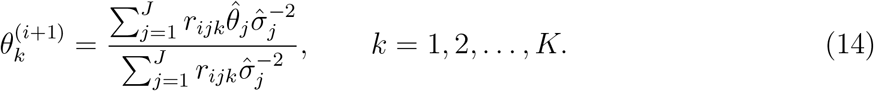

The update equation (14) resembles the inverse-variance weighted (IVW) estimate of the causal effect of the risk factor on the outcome [22, 23]. For a cluster *S* of ratio estimates that target the same causal parameter (that is, the ratio estimates tend to the same causal parameter as the sample size increases), the IVW estimate is the best linear unbiased estimate (BLUE) of this parameter [24]. It is given by

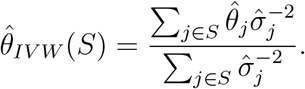

Comparing the above with (14), it follows that the EM update 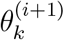 is a re-weighted IVW estimate for the parameter *θ*_*k*_. The weights are multiplied by the responsibilities *r*_*ijk*_ which penalize the influence of observations that are centred away from the current estimate of 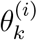 and/or are highly diffuse (that is, 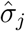 is large). In the large sample limit, as 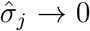 for each *j*, it follows from equation (12) that

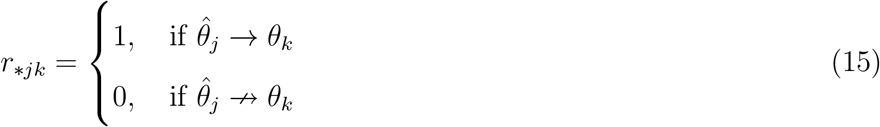

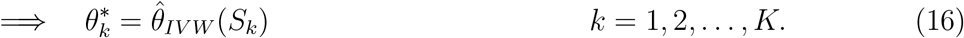

where *r*_\**jk*_ denotes the responsibility of the *k*th component for the *j*th observation computed at an iteration of the EM algorithm in which the MLE is achieved.

The update equations for the mixture proportions *π*_*k*_ are obtained by first modifying equation (11) to account for the constraint that 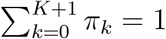 by introducing a Lagrange multiplier, and then maximizing. This is equivalent to solving the following system of equations:

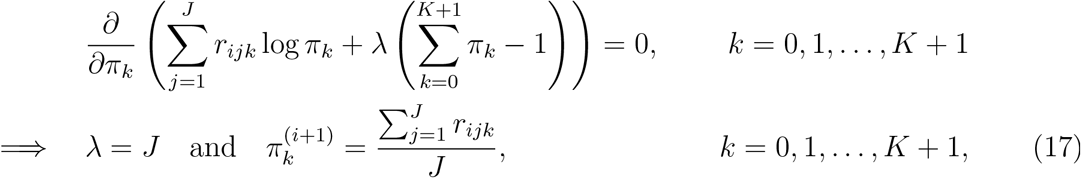

where *λ* denotes the Lagrange multiplier.

### 3.3 Determining the number of clusters

We first calculate the MLEs 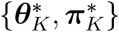 for each value of *K* ∈ {0, 1, …, *J*} possible substantive clusters present in the data. We estimate the number of substantive clusters *K*^*^ by minimizing the Bayesian information criterion (BIC):

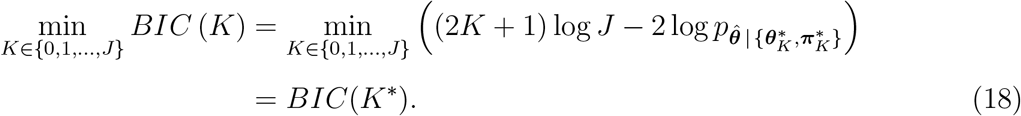

This helps to avoid overparameterization, as the BIC penalizes models which assume the data are generated from larger numbers of underlying clusters.

Pseudocode outlining all steps in the MR-Clust algorithm is given in Algorithm 1. In practice, if *J* is large, then we calculate the MLE and BIC for increasing values of *K* starting at zero, and stop the algorithm once there is evidence that the BIC is increasing monotonically with *K*.

**Algorithm 1:**
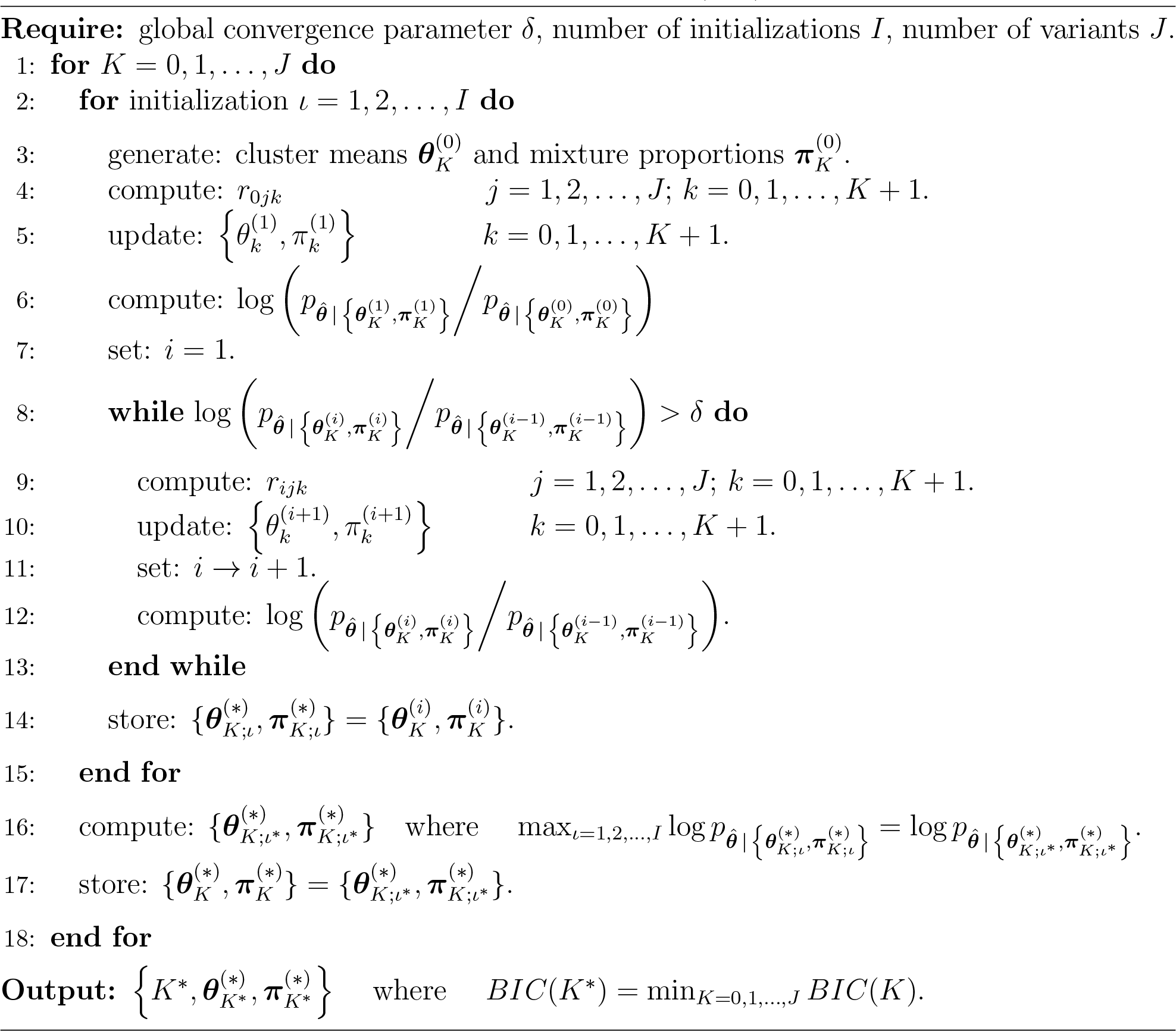
MR-Clust – Expectation Maximization (EM) Algorithm

## 4 Implementation

We perform a simulation study, comparing results from our MR-Clust method to the those obtained using the Mclust method [25]. Mclust is a popular model-based clustering, classification, and density estimation method based on finite normal mixture modelling. Unlike MR-Clust, Mclust does not account for uncertainty in the ratio estimates when assigning observations to clusters. It also does not incorporate null or junk clusters. We show these features help MR-Clust to correctly identify the number of clusters present in the data. We then perform an applied analysis to demonstrate the method in practice.

### 4.1 Simulation: set-up and scenarios

We simulate data on genetic associations with a risk factor 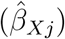 and with an outcome 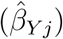 for 50 genetic variants indexed by *j*. These associations imitate coefficient estimates from linear regression of a continuous variable with variance 1 on a single nucleotide polymorphism (SNP). A SNP can be thought of as a binomial random variable taking values 0, 1, 2, representing the number of minor alleles inherited from one’s parents at a particular location of the genetic code.

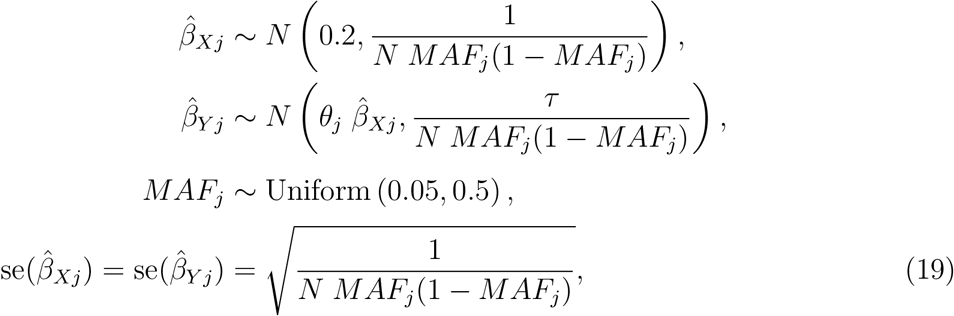

where *N* = 10 000 is the notional sample size in which the genetic associations are estimated, *θ*_*j*_ is the causal effect of variant *j*, *MAF*_*j*_ is the minor allele frequency of variant *j*, and *τ* is an overdispersion parameter.

We consider 4 scenarios, and simulate 1000 datasets in each scenario. In Scenarios 1 and 2, there are no non-null clusters, and *θ*_*j*_ = 0 for all *j*. In Scenarios 3 and 4, there are three non-null clusters *θ*_*j*_ = 0.8 for *j* = 1, …, 10, *θ*_*j*_ = 0.4 for *j* = 11*, …,* 20, *θ*_*j*_ = −0.4 for *j* = 21, …, 30, and one null cluster *θ*_*j*_ = 0 for *j* = 31, …, 50. In Scenarios 1 and 3, we set *τ* = 1 and in Scenarios 2 and 4, we set *τ* = 2. Scenario 1 represents a null scenario, in which all genetic variants should be included in the null cluster. Scenario 2 represents a variance-inflated null scenario, in which all genetic variants should be included in either the null or junk clusters. These scenarios are considered to assess whether the methods find spurious clusters where they do not truly exist. In Scenario 3 and 4, the methods should find three clusters of 10 variants each, and the other 20 variants should be included in either the null (Scenario 3) or the null or junk cluster (Scenario 4).

### 4.2 Simulation results

Results from the simulation study are displayed in Figure 3. We present the Rand index (top panel), which measures the similarity between the true and estimated allocations into clusters [26], and the number of clusters identified by each method (bottom panel). For comparability, for the MR-Clust method we show the number of substantive clusters plus one for the null cluster, as this is the number of clusters in the data as well as the number that the Mclust method should detect. We compare 4 versions of the methods: (A) each variant is assigned to the cluster with the greatest conditional probability (responsibility); (B) variants are only assigned to a cluster if the conditional probability is ≥ 0.8, otherwise they are unassigned; (C) only substantive clusters with at least 4 assigned variants are reported; and (D) the combination of (B) and (D) – only clusters with at least 4 assigned variants at a conditional probability ≥ 0.8 are reported. Version (D) is the strictest version of the method, and is recommended to discourage the reporting of clusters that are evidenced by only a few variants, which therefore may well be spurious. The thresholds of 0.8 for the probability and 4 for the number of variants are arbitrary choice, but gave good performance in the simulation setting.

**Figure 3:**
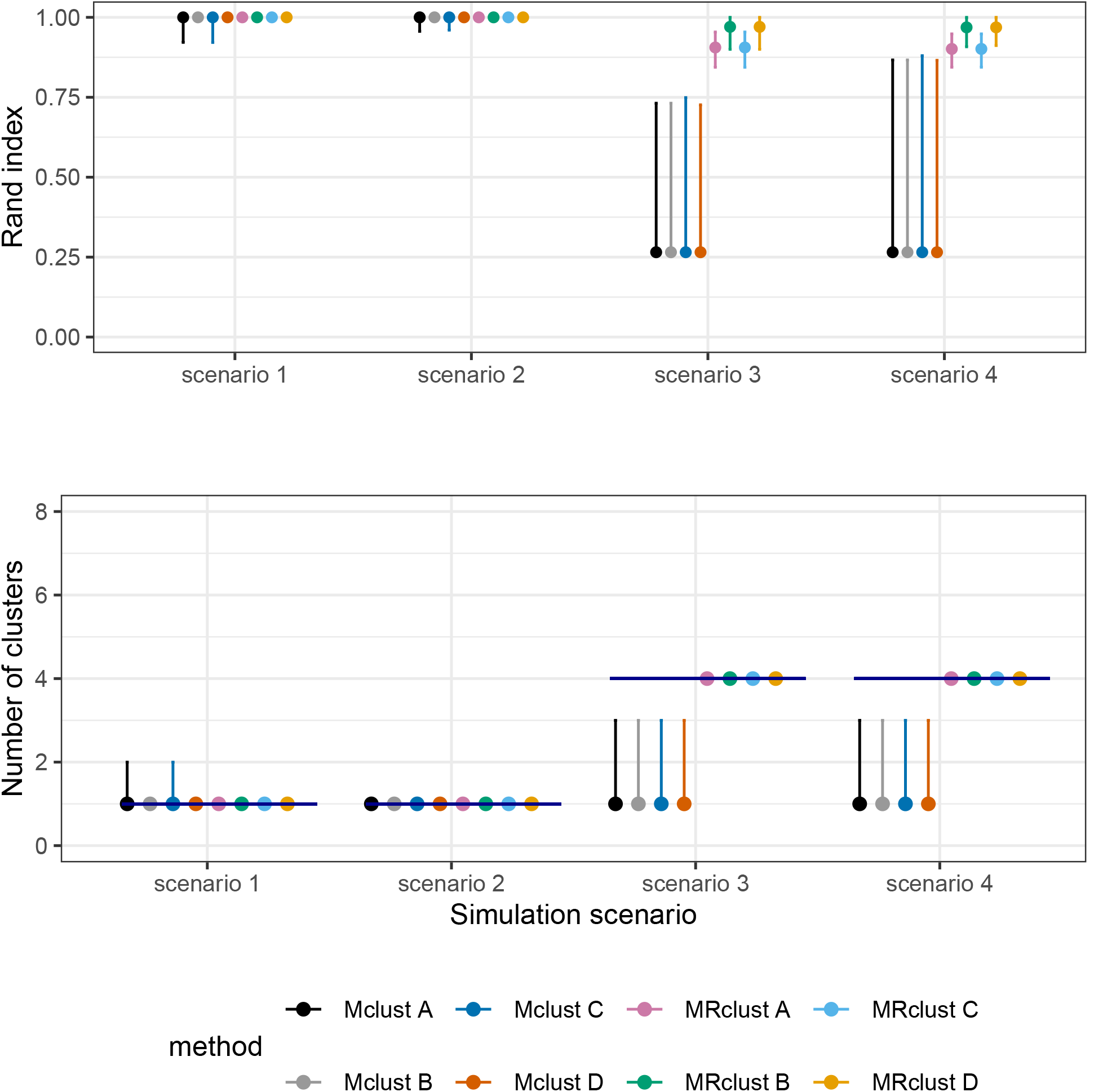
Results from the simulation study for four versions of MR-Clust and Mclust methods under four scenarios for the Rand index (top panel) and the number of clusters identified (bottom panel). Points represent median values across simulated datasets, and vertical bars represent the first and ninth deciles. The horizontal line in the bottom panel represents the true number of clusters in each scenario.

In scenarios 1 and 2, both MR-Clust and Mclust performed similarly across all four versions of the methods (A-D), regularly identifying only a single cluster of variants in the data. In scenarios 3 and 4, results from the two methods differed. MR-Clust continued to perform well in both scenarios, regularly identifying all four clusters in the data. The Rand index was greater for versions (B) and (D), in which clusters with 3 or fewer variants were discounted. Mclust performed poorly in scenarios 3 and 4, typically detecting only one cluster of variants in all versions of the method.

To further illustrate the MR-Clust method, we plot a kernel-weighted density estimate of the distribution of estimated cluster means across the 1000 datasets in Scenario 4 (Figure 4, left panel). On average, MR-Clust identified the correct cluster means at {−0.4, 0, 0.4, 0.8} as well the correct proportions of variants belonging to each cluster. We also plot the value of the log-likelihood at successive iterations of the EM algorithm corresponding to 6 initializations of the parameters for a selected dataset generated under scenario 4 (Figure 4, right panel). The EM algorithm converged to different values of the log-likelihood between the initializations. This indicates some sensitivity of the method to the initial choice of mixture proportions and cluster means, and motivates our use of multiple initializations in the algorithm. We investigated this property across a range of further datasets and simulation scenarios, and usually found negligible differences in MLEs across initializations.

**Figure 4:**
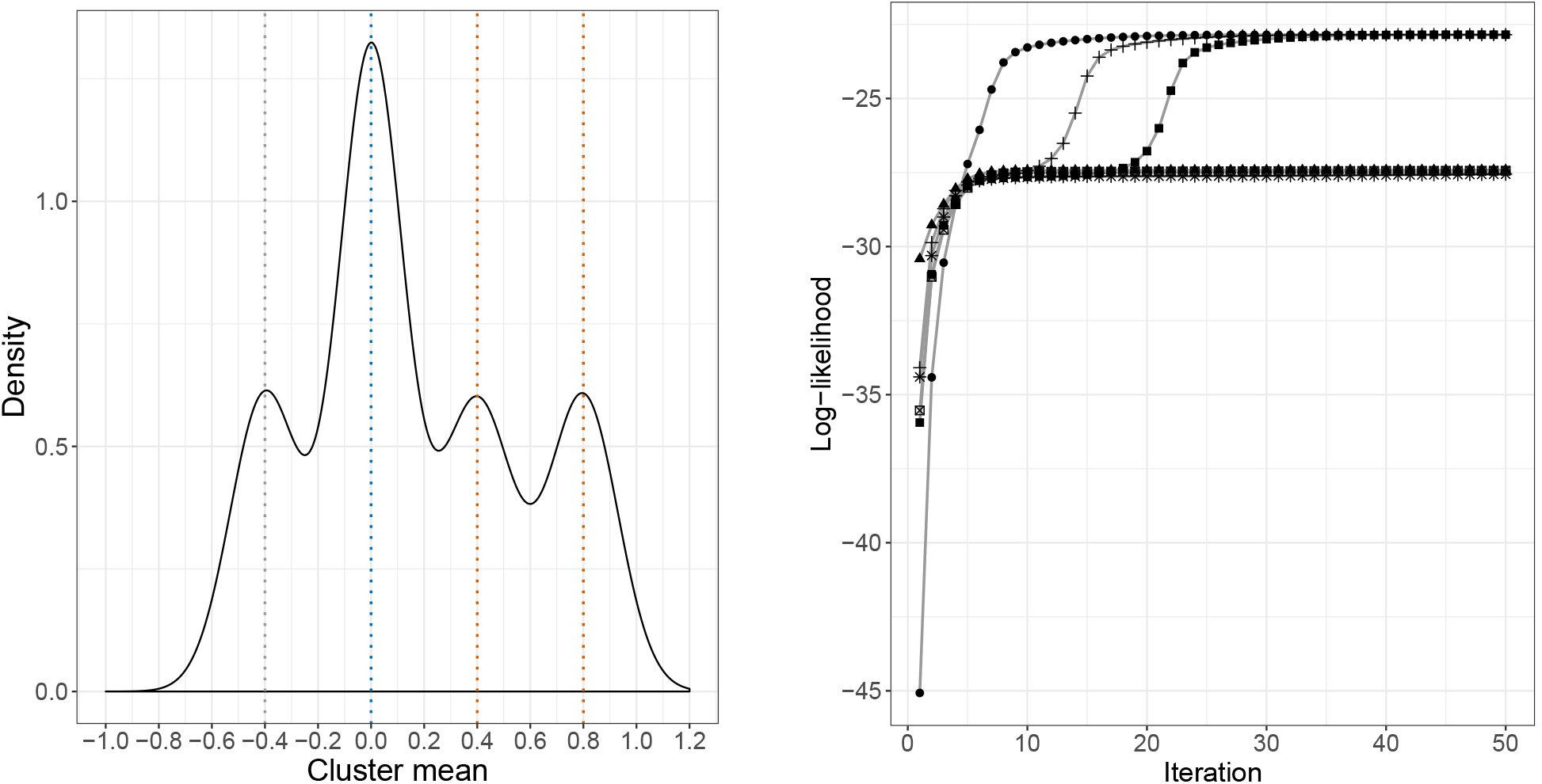
Left panel: Kernel-weighted density plot of cluster means identified by MR-Clust method in simulation scenario 4. Right panel: Convergence to the maximum likelihood estimate across iterations for six initializations for a randomly-chosen dataset from scenario 4 and a fixed number of clusters.

### 4.3 Applied example: blood pressure and coronary artery disease risk

We illustrate our method by considering the relationship between blood pressure and coronary artery disease (CAD) risk. Blood pressure is a heritable trait that is influenced by multiple biological pathways [8, 27]. Elevated blood pressure is considered to be a major risk factor for cardiovascular disease. We assess evidence of clustered heterogeneity in Mendelian randomization analyses for the causal effects of three blood pressure traits on CAD risk: systolic blood pressure (SBP), diastolic blood pressure (DBP) and pulse pressure (PP).

Genetic associations with the blood pressure traits were obtained from the International Consortium for Blood Pressure, and were estimated in 299024 participants of European ancestry [8]. To avoid genetic associations being inflated due to winner’s curse, we only considered genetic variants that had been demonstrated to be associated with a blood pressure trait in a previous genome-wide association study (GWAS) at a genome-wide level of significance (*p* < 5 × 10^−8^). For the analysis of each of the three blood pressure traits, we included all variants additionally associated with the trait under analysis at a p-value threshold of 10^−5^. Variants were all independently distributed (*r*^2^ < 0.01). For SBP, 121 variants were included in the analysis; for DBP, 119 variants; and for PP, 85 variants. Genetic associations with CAD risk were obtained from a meta-analysis of 122733 cases and 424528 controls primarily of European descent from the CARDIoGRAMplusC4D consortium and UK Biobank [28]. We applied the MR-Clust method to assess evidence of clustered heterogeneity for each of the three blood pressure traits on CAD risk separately.

#### Results

Results are displayed in Figure 5. An extract of the results is shown in Table 1, and full results in Supplementary Table A1. Following the simulation study, we present results according to version (A) of the method (all variants assigned to a cluster: top panels) and version (D) (variants assigned to a cluster if conditional probability ≥ 0.8, only clusters with at least 4 variants reported: bottom panels). Although the number of clusters identified varies between SBP and DBP for version (A) of the method, four clusters are reported in version (D) of the method for both traits. This is despite the number and identity of variants varying between the analyses. The largest cluster suggests a positive causal effect of blood pressure on CAD risk. There are also two clusters suggesting a stronger positive causal effect and one suggesting a weak negative effect. For PP, all three substantive clusters in version (D) suggest a positive effect on CAD risk. This suggests the presence of multiple mechanisms by which blood pressure influences CAD risk.

**Table 1.**
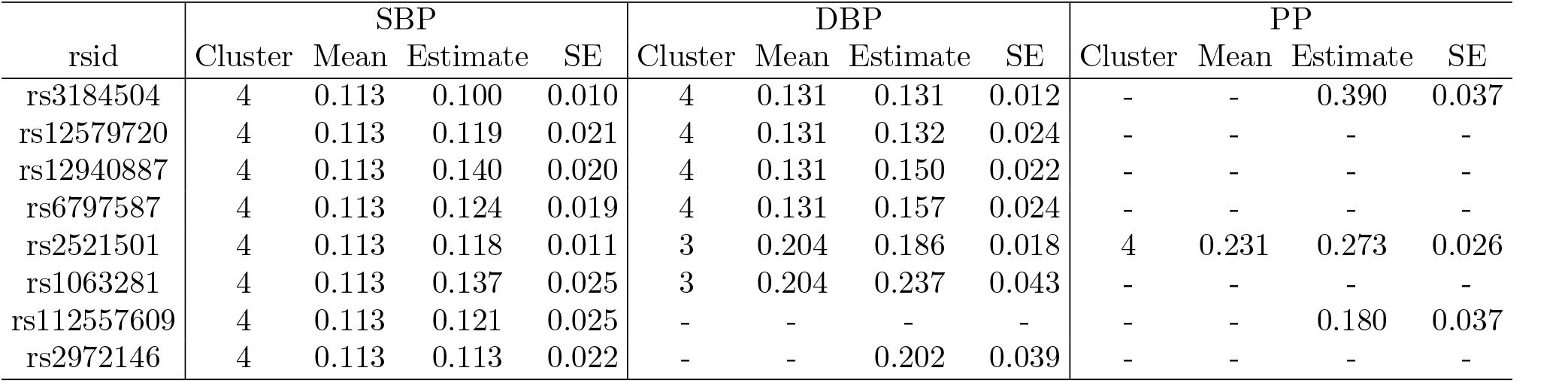
Extract of summary of genetic variants and assignment to clusters in separate analyses for systolic blood pressure (SBP), diastolic blood pressure (DBP), and pulse pressure (PP): cluster number (greatest conditional probability), cluster mean, ratio estimate for that variant, and its standard error. The full results for all 180 variants are in Supplementary Table A1. Dashes either indicate that the variant was not associated with the relevant blood pressure trait at *p* < 10^−5^ (if the estimate is absent), or that the variant was assigned to the null or junk cluster (if the estimate is present).

**Figure 5:**
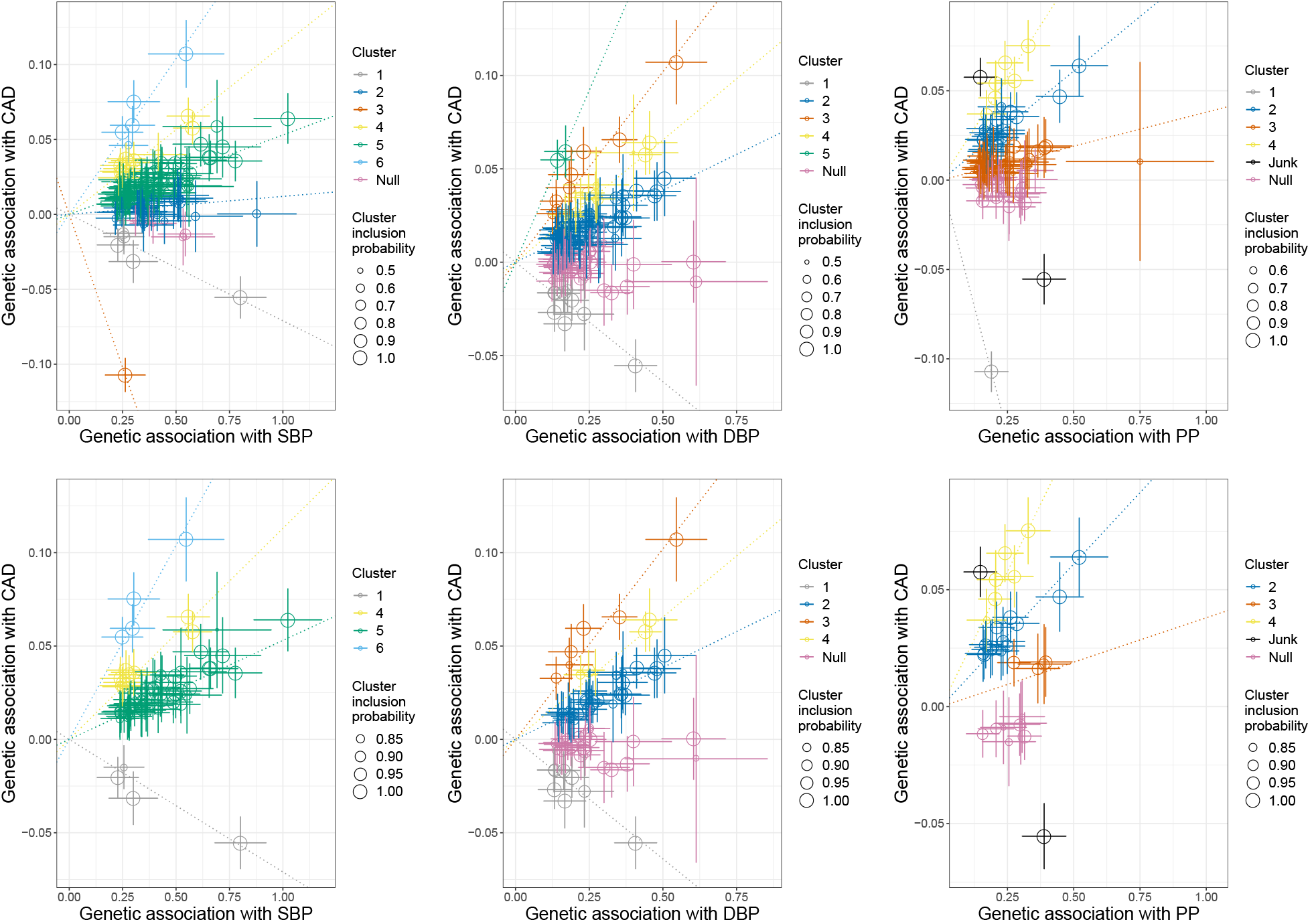
Genetic associations with blood pressure traits (mmHg) and coronary heart disease risk (log odds) per additional blood pressure-increasing allele. Each genetic variant is represented by a point. Error bars are 95% confidence intervals for the genetic associations. Colours represent the clusters, and dotted lines represent the cluster means. Top row: method version (A) – each variant is assigned to the cluster with the greatest conditional probability. Bottom row: method version (D) – variants are only assigned to a cluster if the conditional probability is ≥ 0.8, and clusters are only displayed if at least 4 variants are assigned to the cluster. Left column: systolic blood pressure; middle column: diastolic blood pressure; right column: pulse pressure.

#### Hypothesis-free search for causal mechanism

To demonstrate how clustering can reveal biological mechanisms in the data, we focused on the genetic variants in the clusters for SBP and DBP with a negative effect on CAD risk, and performed a hypothesis-free search of traits that associate with variants in these cluster. We consider this cluster as it is smaller than the two positive clusters, and therefore more plausible that a single mechanism may be driving cluster membership for the majority of variants. In total, 10 genetic variants were assigned to this cluster with a conditional probability ≥ 0.8 in either the SBP or DBP analysis (Supplementary Table A2). We looked up genetic associations in PhenoScanner, a database of genetic associations with traits and diseases [29]. For each trait in turn, we considered whether each variant was associated with that trait at *p* < 10^−5^ and *p* < 10^−8^, and report the true positive rate (the proportion of variants in the cluster associated with the trait) and false positive rate (the proportion of variants not in the cluster associated with the trait). This functionality is built into the *mrclust* software package. In total, we considered 3269 traits, although this includes several repeated or synonymous traits, and blood pressure traits. Also, some traits only had association estimates for a limited number of variants. This makes it difficult to correct for multiple comparisons. We therefore present results under the caveat that no correction has been attempted.

Results are shown in Table 2. The trait “Trunk fat percentage” was associated with 5 out of the 9 variants in the cluster that were present in the dataset (true positive = 0.555), and only 14 out of the 169 variants not in the cluster (false positive = 0.083). Similarly “Impedance of arm right” was associated with 4 out of 9 variants in the cluster (true positive = 0.444), and 15 out of the 169 variants not in the cluster (false positive = 0.089). Impedance is a measure of electrical resistance. It is greater when the body part has a higher fat percentage. At a threshold of *p* < 10^−8^, “Arm fat percentage” was associated with 3 out of 9 variants in the cluster (true positive = 0.333) and only 4 out of the 169 variants not in the cluster (false positive = 0.024).

**Table 2.**
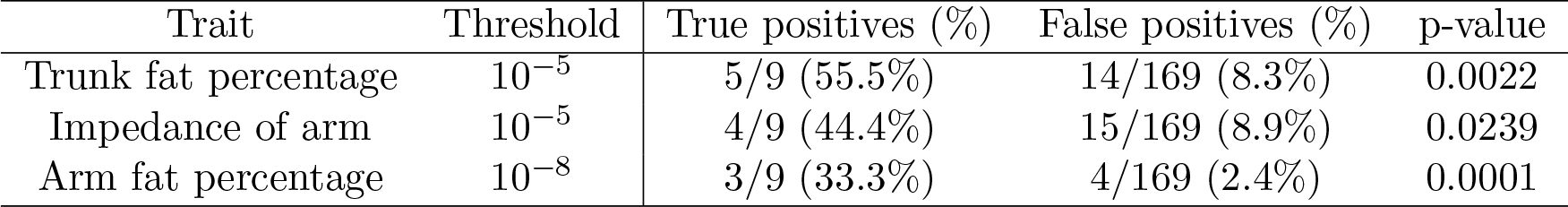
Traits from hypothesis-free search of predictors of cluster membership for cluster with negative causal effect. True positives is the fraction of variants in the cluster that are associated with the trait at a p-value below the threshold. False positives is the fraction of variants not in the cluster that are associated with the trait at a p-value below the threshold. The p-values in the rightmost column are for the hypothesis that association with the trait is independent of cluster membership. P-values are calculated by exact computation from the relevant binomial distribution.

| Trait | Threshold | True positives (%) | False positives (%) | p-value |
| --- | --- | --- | --- | --- |
| Trunk fat percentage | $10^{-5}$ | 5/9 (55.5%) | 14/169 (8.3%) | 0.0022 |
| Impedance of arm | $10^{-5}$ | 4/9 (44.4%) | 15/169 (8.9%) | 0.0239 |
| Arm fat percentage | $10^{-8}$ | 3/9 (33.3%) | 4/169 (2.4%) | 0.0001 |

This suggests that while most biological mechanisms associated with increased blood pressure lead to increased CAD risk, there also may be a biological mechanism associated with decreased blood pressure that leads to increased CAD risk. This mechanism relates to measures of adiposity and fat distribution. However, the directions of association with the adiposity measures were not consistent across variants in the cluster.

## 5 Discussion

In this paper, we have discussed how causal estimates based on different genetic variants in a Mendelian randomization investigation could differ. In particular, we have introduced the notion of clustered heterogeneity, and described how variants that influence the risk factor in different ways could target distinct causal effect parameters. We have introduced the MR-Clust method that detects clusters of variants having similar causal estimates. There are several distinguishing features of this method: it accounts for uncertainty in the causal estimates, and it includes null and junk clusters, so that variants are only included in a substantive cluster if there is strong evidence that they belong to that cluster. We demonstrated the benefits of these features in a simulation study, showing how our method outperforms an existing clustering method that does not have these features. Finally, we illustrated an application of the method to analyse the causal effect of blood pressure on CAD risk, demonstrating the existence of clusters of genetic variants in an empirical example.

In developing this method, our strong concern was to avoid finding spurious clusters of genetic variants that arise due to the chance similarity of causal estimates from different variants. For this reason, we recommend a conservative implementation of the method, only assigning a variant to a cluster if the conditional probability of cluster assignment is ≥ 0.8, and only reporting a cluster if at least 4 variants satisfy this criterion.

Our approach in this paper was to cluster genetic variants based on their causal estimates for a single risk factor and outcome. There are several advantages to this approach. First, there is a natural interpretation of clusters in terms of the causal effect of the risk factor under investigation. Secondly, as the causal estimate is the ratio of the genetic association with the outcome to the genetic association with the risk factor, two variants can appear in the same cluster even if one has weaker associations with the risk factor and outcome, and the other has stronger associations. This is important, as the magnitude of genetic associations is independent of the causal pathway by which it influences the risk factor. Thirdly, as cluster assignment is made on the basis of genetic associations with the risk factor and outcome only, genetic associations with other traits can be used to validate cluster membership, and to explore distinct mechanisms by which the risk factor influences the outcome. If data on genetic associations with multiple traits were used to cluster variants, then the clusters might be more precisely defined, but it would not be possible to determine which traits were driving the division into clusters without further analysis. We have previously demonstrated that a group of variants having similar causal estimates for the effect of HDL-cholesterol on CAD risk also had a distinct pattern of associations with blood cell traits, although without using a formal clustering method [30]. The associations with blood cell traits suggested a causal pathway relating to platelet aggregation.

In order to interpret causal estimates as average causal effects, we made parametric assumptions of linearity and homogeneity. We have discussed these assumptions at length previously [31]. Briefly, the associations of genetic variants with traits are typically small, and so while substantial non-linearity is plausible when considering the causal relationship between a risk factor and outcome across the range of the risk factor distribution, it is less likely when considering the impact of small changes in the average level of the risk factor, as estimated in Mendelian randomization. If the homogeneity assumption is not satisfied, then causal estimates can be interpreted as local average causal effects under the assumption of monotonicity. The monotonicity assumption is generally plausible for genetic variants, as it is difficult to conceive a biological reason why a genetic variant would increase the risk factor in one subset of the population, and decrease it in another. This provides another reason why causal estimates from different genetic variants may differ, as the complier populations corresponding to different genetic variants may differ. However, we believe differences in local average causal effects for different complier populations are unlikely to be substantial in practice. A claim that there are multiple causal pathways from the risk factor to the outcome is more plausible when traits can be found that predict cluster membership, particularly if these traits are potential mediators or moderators of the causal effect of the risk factor.

In conclusion, we have proposed a method in the context of Mendelian randomization that clusters genetic variants associated with a given risk factor according to the variant’s associations with the risk factor and outcome. We have shown theoretically and empirically how the method can help elucidate distinct causal pathways by which the risk factor influences the outcome.

## A Clustered heterogeneity under linearity and homogeneity assumptions

We consider the scenario in which there are linear and homogeneous relationships between genetic variants *G*_1_, …, *G*_*J*_, a risk factor *X*, an outcome *Y*, and a risk factor–outcome confounder *U*. We assume the following linear structural models:

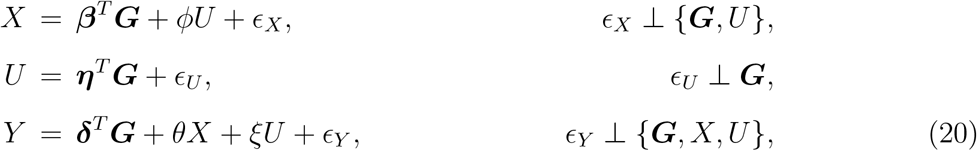

where bold face represents a vector and the epsilon terms represent error in each variable. We assume the *ϕ* and *ξ* are non-zero, and consider ratio estimates for genetic variants with different values of *β*_*j*_, *η*_*j*_ and *δ*_*j*_. The expected value of the ratio estimate (the ratio estimand) using the *j*th variant is:

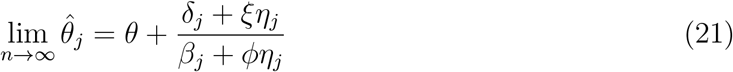

We consider different values of *β*_*j*_, *η*_*j*_ and *δ*_*j*_ that lead to ratio estimands coinciding for variants with different values of these parameters. The two situations in which ratio estimands take the same values for multiple genetic variants with different values of *β*_*j*_, *η*_*j*_ and *δ*_*j*_ are:

1. Genetic variants influence risk factor only (*β*_*j*_ ≠ 0, *η*_*j*_ = 0, *δ*_*j*_ = 0), and
2. Genetic variants influence confounder only (*β*_*j*_ = 0, *η*_*j*_ ≠ 0, *δ*_*j*_ = 0).

This can be seen by exhaustive consideration of all non-zero dimensional subspaces within the overall parameter space. While it is possible for ratio estimands to coincide for other values of *β*_*j*_, *η*_*j*_ and *δ*_*j*_ due to chance, this is vanishingly unlikely.

If we generalize further to a scenario with multiple risk factor–outcome confounders, similar considerations show that ratio estimands will coincide exactly when the genetic variants influence the risk factor only, or one confounder only. This suggests that clustered heterogeneity in the linear and homogeneous scenario corresponds to the situation where genetic variants influence the outcome via a single causal mediator: either the nominated risk factor, or a confounder of the risk factor and outcome. In both situations, there is a common causal pathway from variants in the cluster to the outcome. This corresponds to the diagram of Figure 2.

## B Specification of the junk distribution

To ensure that the distribution of ratio estimates in the junk cluster is near constant across the range of observations from a given sample, whilst also accounting for uncertainty in the ratio estimates 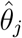 via their standard errors 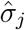, we set the scale parameter *ψ* in the generalised *t*-distribution for the junk cluster to

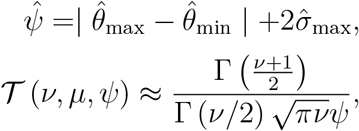

in our applications, where 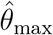 is the maximum of the ratio estimates, 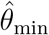 is the minimum of the ratio estimates, and 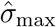 is the maximum of the standard errors of the ratio estimates. This means that the density of the junk cluster will vary between samples, automatically accounting for differences in the range and precision of the ratio estimates. The junk cluster density is a proper density that is approximately uniform across the range of plausible values of the ratio estimates.

**Table A1:**
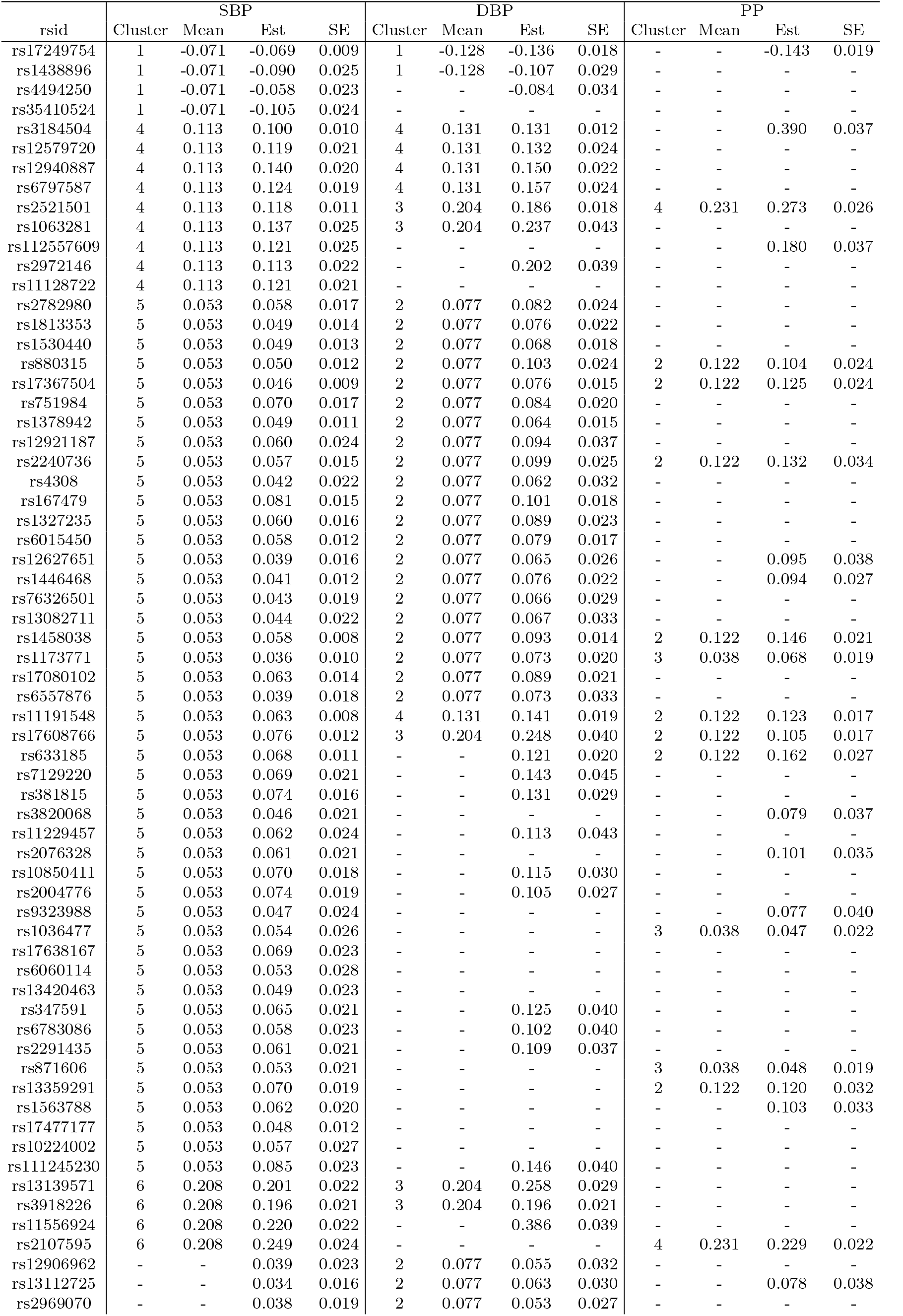
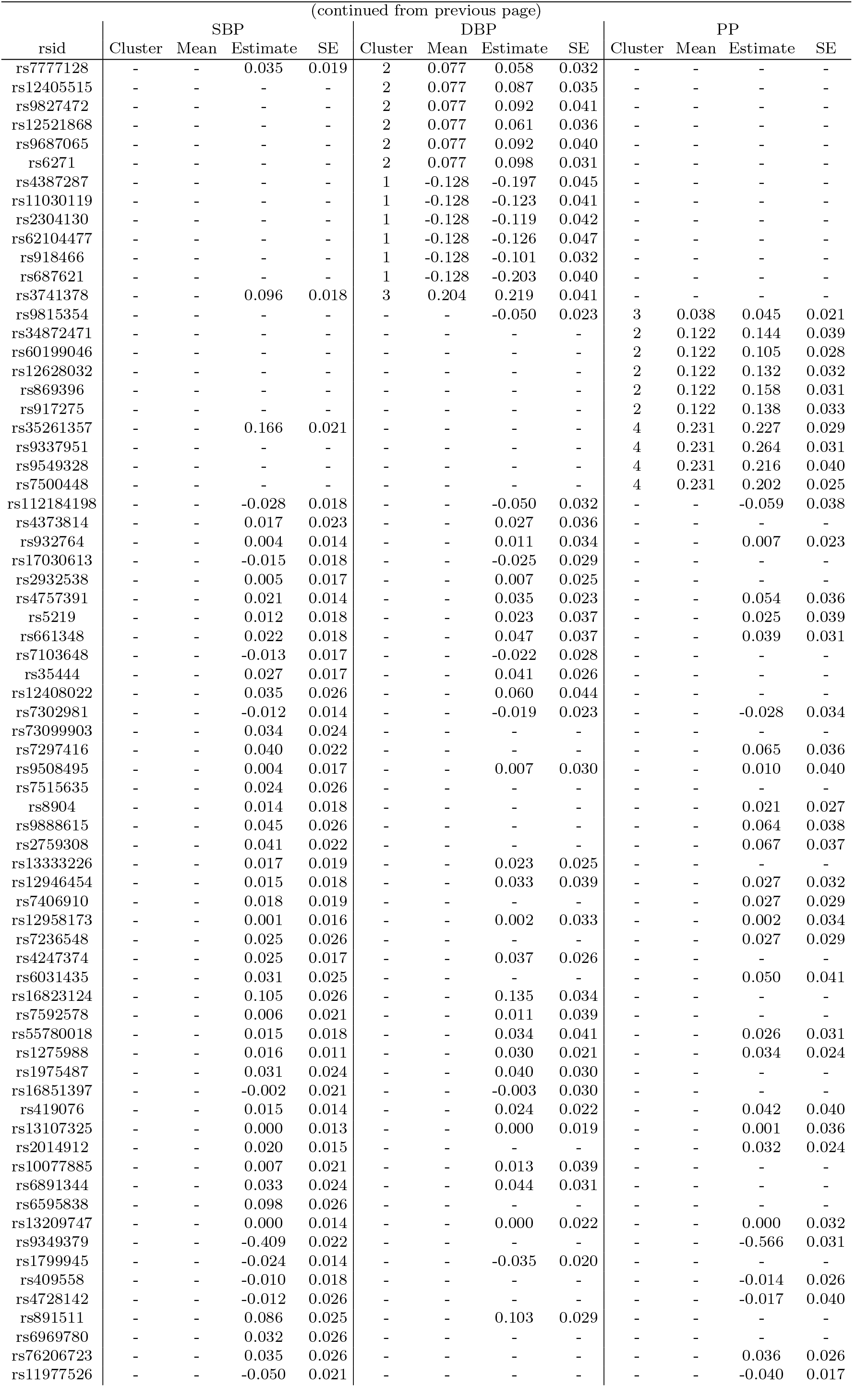
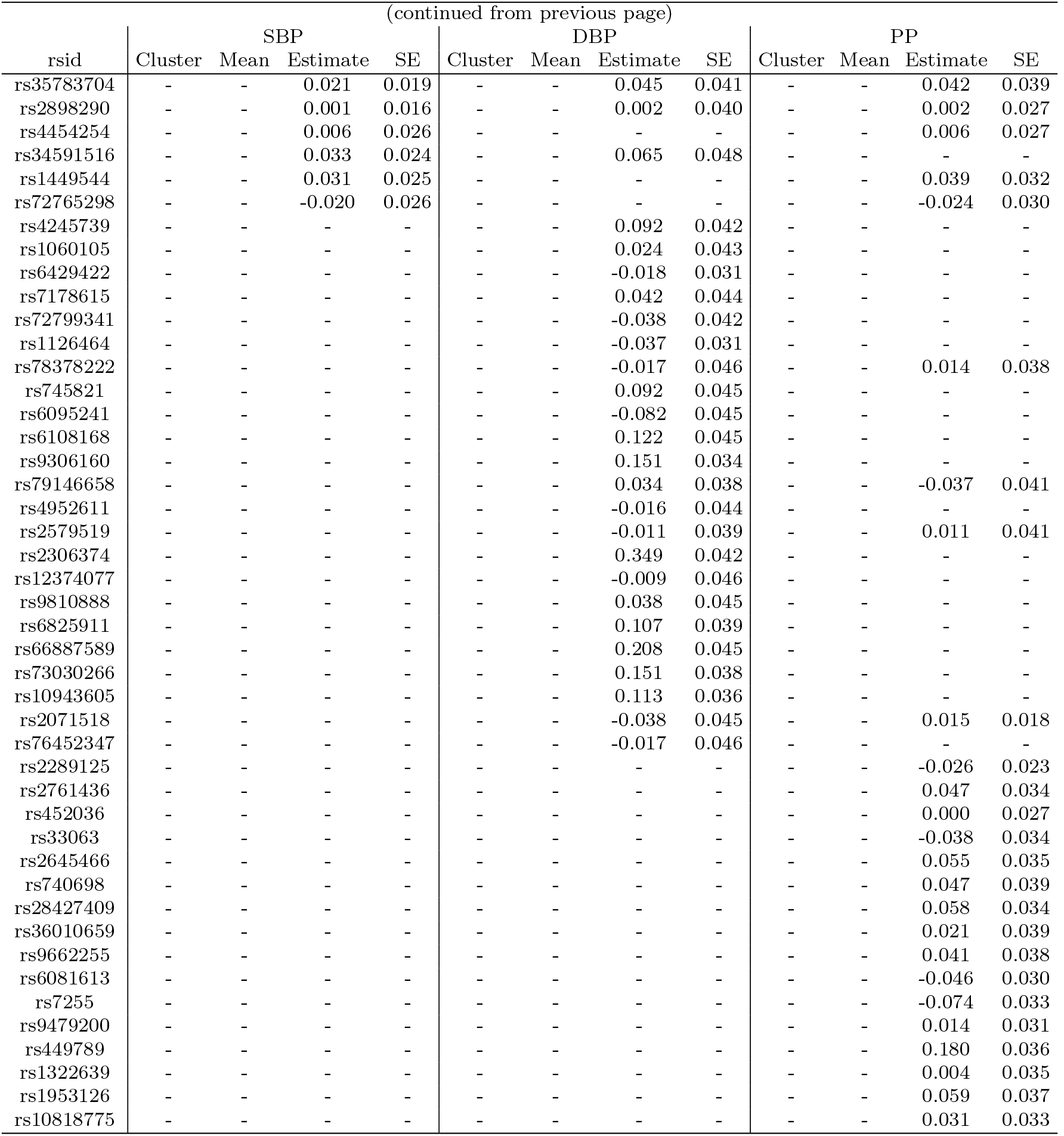
Summary of genetic variants and assignment to clusters in separate analyses for systolic blood pressure (SBP), diastolic blood pressure (DBP), and pulse pressure (PP): cluster number (based on highest conditional probability), cluster mean, ratio estimate (Estimate) for that variant, and its standard error (SE).

**Table A2:**
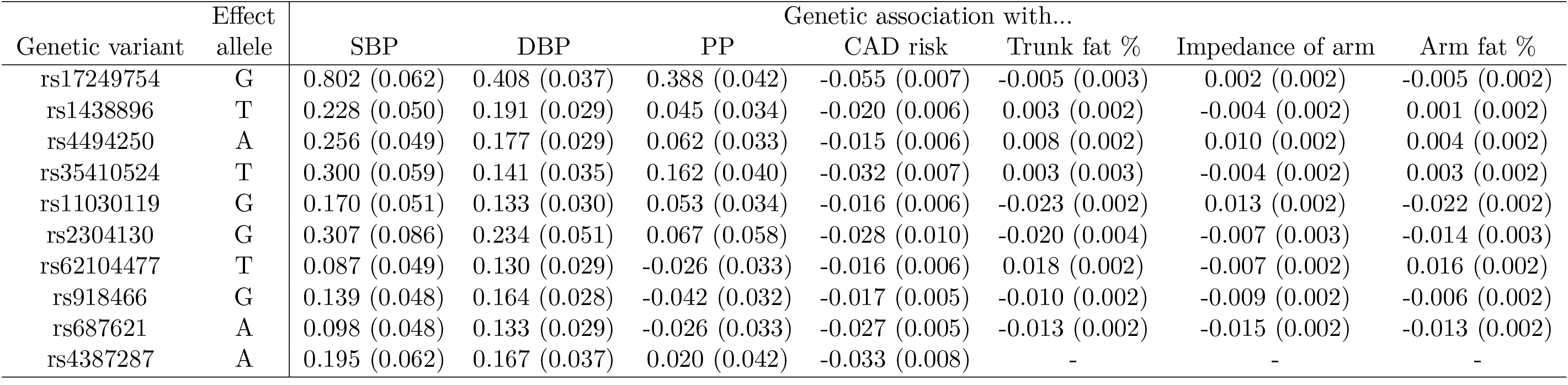
Genetic associations (beta-coefficients and standard errors) for variants in the cluster with negative causal effect of blood pressure on coronary artery disease risk. Genetic associations with blood pressure (mmHg) were estimated in 299 024 participants of European ancestry from the International Consortium for Blood Pressure. Genetic associations with coronary artery disease (CAD) risk (log odds ratios) were estimated in 122 733 cases and 424 528 controls primarily of European descent from the CARDIoGRAMplusC4D consortium and UK Biobank. Genetic associations with the adiposity traits (SD units) were estimated in 337 199 participants of European descent from UK Biobank (Ben Neale estimates). Associations in UK Biobank for rs4387287 were not available.

