## Supplementary material for "MR-Clust: Clustering of genetic variants in Mendelian randomization with similar causal estimates"

### A Clustered heterogeneity under linearity and homogeneity assumptions

We consider the scenario in which there are linear and homogeneous relationships between genetic variants $G_1, \ldots, G_J$, a risk factor $X$, an outcome $Y$, and a risk factor–outcome confounder $U$. We assume the following linear structural models:

$$\begin{aligned} X &= \boldsymbol{\beta}^T \boldsymbol{G} + \phi U + \epsilon_X, & \epsilon_X &\perp \{\boldsymbol{G}, U\}, \\ U &= \boldsymbol{\eta}^T \boldsymbol{G} + \epsilon_U, & \epsilon_U &\perp \boldsymbol{G}, \\ Y &= \boldsymbol{\delta}^T \boldsymbol{G} + \theta X + \xi U + \epsilon_Y, & \epsilon_Y &\perp \{\boldsymbol{G}, X, U\}, \end{aligned} \tag{20}$$

where bold face represents a vector and the epsilon terms represent error in each variable. We assume the $\phi$ and $\xi$ are non-zero, and consider ratio estimates for genetic variants with different values of $\beta_j$, $\eta_j$ and $\delta_j$. The expected value of the ratio estimate (the ratio estimand) using the $j$th variant is:

$$\lim_{n\to\infty} \hat{\theta}_j = \theta + \frac{\delta_j + \xi\eta_j}{\beta_j + \phi\eta_j} \tag{21}$$

We consider different values of $\beta_j$, $\eta_j$ and $\delta_j$ that lead to ratio estimands coinciding for variants with different values of these parameters. The two situations in which ratio estimands take the same values for multiple genetic variants with different values of $\beta_j$, $\eta_j$ and $\delta_j$ are:

1. Genetic variants influence risk factor only ($\beta_j \neq 0$, $\eta_j = 0$, $\delta_j = 0$), and

2. Genetic variants influence confounder only ($\beta_j = 0$, $\eta_j \neq 0$, $\delta_j = 0$).

This can be seen by exhaustive consideration of all non-zero dimensional subspaces within the overall parameter space. While it is possible for ratio estimands to coincide for other values of $\beta_j$, $\eta_j$ and $\delta_j$ due to chance, this is vanishingly unlikely.

If we generalize further to a scenario with multiple risk factor–outcome confounders, similar considerations show that ratio estimands will coincide exactly when the genetic variants influence the risk factor only, or one confounder only. This suggests that clustered heterogeneity in the linear and homogeneous scenario corresponds to the situation where genetic variants influence the outcome via a single causal mediator: either the nominated risk factor, or a confounder of the risk factor and outcome. In both situations, there is a common causal pathway from variants in the cluster to the outcome. This corresponds to the diagram of Figure 2.

#### B Specification of the junk distribution

To ensure that the distribution of ratio estimates in the junk cluster is near constant across the range of observations from a given sample, whilst also accounting for uncertainty in the ratio estimates $\hat{\theta}_j$ via their standard errors $\hat{\sigma}_j$, we set the scale parameter $\psi$ in the generalised $t$-distribution for the junk cluster to

$$\hat{\psi} = \mid \hat{\theta}_{\text{max}} - \hat{\theta}_{\text{min}} \mid + 2\hat{\sigma}_{\text{max}},$$
$$\mathcal{T}\left(\nu, \mu, \psi\right) \approx \frac{\Gamma\left(\frac{\nu+1}{2}\right)}{\Gamma\left(\nu/2\right)\sqrt{\pi\nu}\psi},$$

in our applications, where $\hat{\theta}_{\text{max}}$ is the maximum of the ratio estimates, $\hat{\theta}_{\text{min}}$ is the minimum of the ratio estimates, and $\hat{\sigma}_{\text{max}}$ is the maximum of the standard errors of the ratio estimates. This means that the density of the junk cluster will vary between samples, automatically accounting for differences in the range and precision of the ratio estimates. The junk cluster density is a proper density that is approximately uniform across the range of plausible values of the ratio estimates.

Table A1: Summary of genetic variants and assignment to clusters in separate analyses for systolic blood pressure (SBP), diastolic blood pressure (DBP), and pulse pressure (PP): cluster number (based on highest conditional probability), cluster mean, ratio estimate (Estimate) for that variant, and its standard error (SE).

| | SBP | | | | DBP | | | | PP | | | |
|---|---|---|---|---|---|---|---|---|---|---|---|---|
| rsid | Cluster | Mean | Est | SE | Cluster | Mean | Est | SE | Cluster | Mean | Est | SE |
| rs17249754 | 1 | -0.071 | -0.069 | 0.009 | 1 | -0.128 | -0.136 | 0.018 | - | - | -0.143 | 0.019 |
| rs1438896 | 1 | -0.071 | -0.090 | 0.025 | 1 | -0.128 | -0.107 | 0.029 | - | - | - | - |
| rs4494250 | 1 | -0.071 | -0.058 | 0.023 | - | - | -0.084 | 0.034 | - | - | - | - |
| rs35410524 | 1 | -0.071 | -0.105 | 0.024 | - | - | - | - | - | - | - | - |
| rs3184504 | 4 | 0.113 | 0.100 | 0.010 | 4 | 0.131 | 0.131 | 0.012 | - | - | 0.390 | 0.037 |
| rs12579720 | 4 | 0.113 | 0.119 | 0.021 | 4 | 0.131 | 0.132 | 0.024 | - | - | - | - |
| rs12940887 | 4 | 0.113 | 0.140 | 0.020 | 4 | 0.131 | 0.150 | 0.022 | - | - | - | - |
| rs6797587 | 4 | 0.113 | 0.124 | 0.019 | 4 | 0.131 | 0.157 | 0.024 | - | - | - | - |
| rs2521501 | 4 | 0.113 | 0.118 | 0.011 | 3 | 0.204 | 0.186 | 0.018 | 4 | 0.231 | 0.273 | 0.026 |
| rs1063281 | 4 | 0.113 | 0.137 | 0.025 | 3 | 0.204 | 0.237 | 0.043 | - | - | - | - |
| rs112557609 | 4 | 0.113 | 0.121 | 0.025 | - | - | - | - | - | - | 0.180 | 0.037 |
| rs2972146 | 4 | 0.113 | 0.113 | 0.022 | - | - | 0.202 | 0.039 | - | - | - | - |
| rs11128722 | 4 | 0.113 | 0.121 | 0.021 | - | - | - | - | - | - | - | - |
| rs2782980 | 5 | 0.053 | 0.058 | 0.017 | 2 | 0.077 | 0.082 | 0.024 | - | - | - | - |
| rs1813353 | 5 | 0.053 | 0.049 | 0.014 | 2 | 0.077 | 0.076 | 0.022 | - | - | - | - |
| rs1530440 | 5 | 0.053 | 0.049 | 0.013 | 2 | 0.077 | 0.068 | 0.018 | - | - | - | - |
| rs880315 | 5 | 0.053 | 0.050 | 0.012 | 2 | 0.077 | 0.103 | 0.024 | 2 | 0.122 | 0.104 | 0.024 |
| rs17367504 | 5 | 0.053 | 0.046 | 0.009 | 2 | 0.077 | 0.076 | 0.015 | 2 | 0.122 | 0.125 | 0.024 |
| rs751984 | 5 | 0.053 | 0.070 | 0.017 | 2 | 0.077 | 0.084 | 0.020 | - | - | - | - |
| rs1378942 | 5 | 0.053 | 0.049 | 0.011 | 2 | 0.077 | 0.064 | 0.015 | - | - | - | - |
| rs12921187 | 5 | 0.053 | 0.060 | 0.024 | 2 | 0.077 | 0.094 | 0.037 | - | - | - | - |
| rs2240736 | 5 | 0.053 | 0.057 | 0.015 | 2 | 0.077 | 0.099 | 0.025 | 2 | 0.122 | 0.132 | 0.034 |
| rs4308 | 5 | 0.053 | 0.042 | 0.022 | 2 | 0.077 | 0.062 | 0.032 | - | - | - | - |
| rs167479 | 5 | 0.053 | 0.081 | 0.015 | 2 | 0.077 | 0.101 | 0.018 | - | - | - | - |
| rs1327235 | 5 | 0.053 | 0.060 | 0.016 | 2 | 0.077 | 0.089 | 0.023 | - | - | - | - |
| rs6015450 | 5 | 0.053 | 0.058 | 0.012 | 2 | 0.077 | 0.079 | 0.017 | - | - | - | - |
| rs12627651 | 5 | 0.053 | 0.039 | 0.016 | 2 | 0.077 | 0.065 | 0.026 | - | - | 0.095 | 0.038 |
| rs1446468 | 5 | 0.053 | 0.041 | 0.012 | 2 | 0.077 | 0.076 | 0.022 | - | - | 0.094 | 0.027 |
| rs76326501 | 5 | 0.053 | 0.043 | 0.019 | 2 | 0.077 | 0.066 | 0.029 | - | - | - | - |
| rs13082711 | 5 | 0.053 | 0.044 | 0.022 | 2 | 0.077 | 0.067 | 0.033 | - | - | - | - |
| rs1458038 | 5 | 0.053 | 0.058 | 0.008 | 2 | 0.077 | 0.093 | 0.014 | 2 | 0.122 | 0.146 | 0.021 |
| rs1173771 | 5 | 0.053 | 0.036 | 0.010 | 2 | 0.077 | 0.073 | 0.020 | 3 | 0.038 | 0.068 | 0.019 |
| rs17080102 | 5 | 0.053 | 0.063 | 0.014 | 2 | 0.077 | 0.089 | 0.021 | - | - | - | - |
| rs6557876 | 5 | 0.053 | 0.039 | 0.018 | 2 | 0.077 | 0.073 | 0.033 | - | - | - | - |
| rs11191548 | 5 | 0.053 | 0.063 | 0.008 | 4 | 0.131 | 0.141 | 0.019 | 2 | 0.122 | 0.123 | 0.017 |
| rs17608766 | 5 | 0.053 | 0.076 | 0.012 | 3 | 0.204 | 0.248 | 0.040 | 2 | 0.122 | 0.105 | 0.017 |
| rs633185 | 5 | 0.053 | 0.068 | 0.011 | - | - | 0.121 | 0.020 | 2 | 0.122 | 0.162 | 0.027 |
| rs7129220 | 5 | 0.053 | 0.069 | 0.021 | - | - | 0.143 | 0.045 | - | - | - | - |
| rs381815 | 5 | 0.053 | 0.074 | 0.016 | - | - | 0.131 | 0.029 | - | - | - | - |
| rs3820068 | 5 | 0.053 | 0.046 | 0.021 | - | - | - | - | - | - | 0.079 | 0.037 |
| rs11229457 | 5 | 0.053 | 0.062 | 0.024 | - | - | 0.113 | 0.043 | - | - | - | - |
| rs2076328 | 5 | 0.053 | 0.061 | 0.021 | - | - | - | - | - | - | 0.101 | 0.035 |
| rs10850411 | 5 | 0.053 | 0.070 | 0.018 | - | - | 0.115 | 0.030 | - | - | - | - |
| rs2004776 | 5 | 0.053 | 0.074 | 0.019 | - | - | 0.105 | 0.027 | - | - | - | - |
| rs9323988 | 5 | 0.053 | 0.047 | 0.024 | - | - | - | - | - | - | 0.077 | 0.040 |
| rs1036477 | 5 | 0.053 | 0.054 | 0.026 | - | - | - | - | 3 | 0.038 | 0.047 | 0.022 |
| rs17638167 | 5 | 0.053 | 0.069 | 0.023 | - | - | - | - | - | - | - | - |
| rs6060114 | 5 | 0.053 | 0.053 | 0.028 | - | - | - | - | - | - | - | - |
| rs13420463 | 5 | 0.053 | 0.049 | 0.023 | - | - | - | - | - | - | - | - |
| rs347591 | 5 | 0.053 | 0.065 | 0.021 | - | - | 0.125 | 0.040 | - | - | - | - |
| rs6783086 | 5 | 0.053 | 0.058 | 0.023 | - | - | 0.102 | 0.040 | - | - | - | - |
| rs2291435 | 5 | 0.053 | 0.061 | 0.021 | - | - | 0.109 | 0.037 | - | - | - | - |
| rs871606 | 5 | 0.053 | 0.053 | 0.021 | - | - | - | - | 3 | 0.038 | 0.048 | 0.019 |
| rs13359291 | 5 | 0.053 | 0.070 | 0.019 | - | - | - | - | 2 | 0.122 | 0.120 | 0.032 |
| rs1563788 | 5 | 0.053 | 0.062 | 0.020 | - | - | - | - | - | - | 0.103 | 0.033 |
| rs17477177 | 5 | 0.053 | 0.048 | 0.012 | - | - | - | - | - | - | - | - |
| rs10224002 | 5 | 0.053 | 0.057 | 0.027 | - | - | - | - | - | - | - | - |
| rs111245230 | 5 | 0.053 | 0.085 | 0.023 | - | - | 0.146 | 0.040 | - | - | - | - |
| rs13139571 | 6 | 0.208 | 0.201 | 0.022 | 3 | 0.204 | 0.258 | 0.029 | - | - | - | - |
| rs3918226 | 6 | 0.208 | 0.196 | 0.021 | 3 | 0.204 | 0.196 | 0.021 | - | - | - | - |
| rs11556924 | 6 | 0.208 | 0.220 | 0.022 | - | - | 0.386 | 0.039 | - | - | - | - |
| rs2107595 | 6 | 0.208 | 0.249 | 0.024 | - | - | - | - | 4 | 0.231 | 0.229 | 0.022 |
| rs12906962 | - | - | 0.039 | 0.023 | 2 | 0.077 | 0.055 | 0.032 | - | - | - | - |
| rs13112725 | - | - | 0.034 | 0.016 | 2 | 0.077 | 0.063 | 0.030 | - | - | 0.078 | 0.038 |
| rs2969070 | - | - | 0.038 | 0.019 | 2 | 0.077 | 0.053 | 0.027 | - | - | - | - |

(continued from previous page)

| | SBP | | | | DBP | | | | PP | | | |
|---|---|---|---|---|---|---|---|---|---|---|---|---|
| rsid | Cluster | Mean | Estimate | SE | Cluster | Mean | Estimate | SE | Cluster | Mean | Estimate | SE |
| rs7777128 | - | - | 0.035 | 0.019 | 2 | 0.077 | 0.058 | 0.032 | - | - | - | - |
| rs12405515 | - | - | - | - | 2 | 0.077 | 0.087 | 0.035 | - | - | - | - |
| rs9827472 | - | - | - | - | 2 | 0.077 | 0.092 | 0.041 | - | - | - | - |
| rs12521868 | - | - | - | - | 2 | 0.077 | 0.061 | 0.036 | - | - | - | - |
| rs9687065 | - | - | - | - | 2 | 0.077 | 0.092 | 0.040 | - | - | - | - |
| rs6271 | - | - | - | - | 2 | 0.077 | 0.098 | 0.031 | - | - | - | - |
| rs4387287 | - | - | - | - | 1 | -0.128 | -0.197 | 0.045 | - | - | - | - |
| rs11030119 | - | - | - | - | 1 | -0.128 | -0.123 | 0.041 | - | - | - | - |
| rs2304130 | - | - | - | - | 1 | -0.128 | -0.119 | 0.042 | - | - | - | - |
| rs62104477 | - | - | - | - | 1 | -0.128 | -0.126 | 0.047 | - | - | - | - |
| rs918466 | - | - | - | - | 1 | -0.128 | -0.101 | 0.032 | - | - | - | - |
| rs687621 | - | - | - | - | 1 | -0.128 | -0.203 | 0.040 | - | - | - | - |
| rs3741378 | - | - | 0.096 | 0.018 | 3 | 0.204 | 0.219 | 0.041 | - | - | - | - |
| rs9815354 | - | - | - | - | - | - | -0.050 | 0.023 | 3 | 0.038 | 0.045 | 0.021 |
| rs34872471 | - | - | - | - | - | - | - | - | 2 | 0.122 | 0.144 | 0.039 |
| rs60199046 | - | - | - | - | - | - | - | - | 2 | 0.122 | 0.105 | 0.028 |
| rs12628032 | - | - | - | - | - | - | - | - | 2 | 0.122 | 0.132 | 0.032 |
| rs869396 | - | - | - | - | - | - | - | - | 2 | 0.122 | 0.158 | 0.031 |
| rs917275 | - | - | - | - | - | - | - | - | 2 | 0.122 | 0.138 | 0.033 |
| rs35261357 | - | - | 0.166 | 0.021 | - | - | - | - | 4 | 0.231 | 0.227 | 0.029 |
| rs9337951 | - | - | - | - | - | - | - | - | 4 | 0.231 | 0.264 | 0.031 |
| rs9549328 | - | - | - | - | - | - | - | - | 4 | 0.231 | 0.216 | 0.040 |
| rs7500448 | - | - | - | - | - | - | - | - | 4 | 0.231 | 0.202 | 0.025 |
| rs112184198 | - | - | -0.028 | 0.018 | - | - | -0.050 | 0.032 | - | - | -0.059 | 0.038 |
| rs4373814 | - | - | 0.017 | 0.023 | - | - | 0.027 | 0.036 | - | - | - | - |
| rs932764 | - | - | 0.004 | 0.014 | - | - | 0.011 | 0.034 | - | - | 0.007 | 0.023 |
| rs17030613 | - | - | -0.015 | 0.018 | - | - | -0.025 | 0.029 | - | - | - | - |
| rs2932538 | - | - | 0.005 | 0.017 | - | - | 0.007 | 0.025 | - | - | - | - |
| rs4757391 | - | - | 0.021 | 0.014 | - | - | 0.035 | 0.023 | - | - | 0.054 | 0.036 |
| rs5219 | - | - | 0.012 | 0.018 | - | - | 0.023 | 0.037 | - | - | 0.025 | 0.039 |
| rs661348 | - | - | 0.022 | 0.018 | - | - | 0.047 | 0.037 | - | - | 0.039 | 0.031 |
| rs7103648 | - | - | -0.013 | 0.017 | - | - | -0.022 | 0.028 | - | - | - | - |
| rs35444 | - | - | 0.027 | 0.017 | - | - | 0.041 | 0.026 | - | - | - | - |
| rs12408022 | - | - | 0.035 | 0.026 | - | - | 0.060 | 0.044 | - | - | - | - |
| rs7302981 | - | - | -0.012 | 0.014 | - | - | -0.019 | 0.023 | - | - | -0.028 | 0.034 |
| rs73099903 | - | - | 0.034 | 0.024 | - | - | - | - | - | - | - | - |
| rs7297416 | - | - | 0.040 | 0.022 | - | - | - | - | - | - | 0.065 | 0.036 |
| rs9508495 | - | - | 0.004 | 0.017 | - | - | 0.007 | 0.030 | - | - | 0.010 | 0.040 |
| rs7515635 | - | - | 0.024 | 0.026 | - | - | - | - | - | - | - | - |
| rs8904 | - | - | 0.014 | 0.018 | - | - | - | - | - | - | 0.021 | 0.027 |
| rs9888615 | - | - | 0.045 | 0.026 | - | - | - | - | - | - | 0.064 | 0.038 |
| rs2759308 | - | - | 0.041 | 0.022 | - | - | - | - | - | - | 0.067 | 0.037 |
| rs13333226 | - | - | 0.017 | 0.019 | - | - | 0.023 | 0.025 | - | - | - | - |
| rs12946454 | - | - | 0.015 | 0.018 | - | - | 0.033 | 0.039 | - | - | 0.027 | 0.032 |
| rs7406910 | - | - | 0.018 | 0.019 | - | - | - | - | - | - | 0.027 | 0.029 |
| rs12958173 | - | - | 0.001 | 0.016 | - | - | 0.002 | 0.033 | - | - | 0.002 | 0.034 |
| rs7236548 | - | - | 0.025 | 0.026 | - | - | - | - | - | - | 0.027 | 0.029 |
| rs4247374 | - | - | 0.025 | 0.017 | - | - | 0.037 | 0.026 | - | - | - | - |
| rs6031435 | - | - | 0.031 | 0.025 | - | - | - | - | - | - | 0.050 | 0.041 |
| rs16823124 | - | - | 0.105 | 0.026 | - | - | 0.135 | 0.034 | - | - | - | - |
| rs7592578 | - | - | 0.006 | 0.021 | - | - | 0.011 | 0.039 | - | - | - | - |
| rs55780018 | - | - | 0.015 | 0.018 | - | - | 0.034 | 0.041 | - | - | 0.026 | 0.031 |
| rs1275988 | - | - | 0.016 | 0.011 | - | - | 0.030 | 0.021 | - | - | 0.034 | 0.024 |
| rs1975487 | - | - | 0.031 | 0.024 | - | - | 0.040 | 0.030 | - | - | - | - |
| rs16851397 | - | - | -0.002 | 0.021 | - | - | -0.003 | 0.030 | - | - | - | - |
| rs419076 | - | - | 0.015 | 0.014 | - | - | 0.024 | 0.022 | - | - | 0.042 | 0.040 |
| rs13107325 | - | - | 0.000 | 0.013 | - | - | 0.000 | 0.019 | - | - | 0.001 | 0.036 |
| rs2014912 | - | - | 0.020 | 0.015 | - | - | - | - | - | - | 0.032 | 0.024 |
| rs10077885 | - | - | 0.007 | 0.021 | - | - | 0.013 | 0.039 | - | - | - | - |
| rs6891344 | - | - | 0.033 | 0.024 | - | - | 0.044 | 0.031 | - | - | - | - |
| rs6595838 | - | - | 0.098 | 0.026 | - | - | - | - | - | - | - | - |
| rs13209747 | - | - | 0.000 | 0.014 | - | - | 0.000 | 0.022 | - | - | 0.000 | 0.032 |
| rs9349379 | - | - | -0.409 | 0.022 | - | - | - | - | - | - | -0.566 | 0.031 |
| rs1799945 | - | - | -0.024 | 0.014 | - | - | -0.035 | 0.020 | - | - | - | - |
| rs409558 | - | - | -0.010 | 0.018 | - | - | - | - | - | - | -0.014 | 0.026 |
| rs4728142 | - | - | -0.012 | 0.026 | - | - | - | - | - | - | -0.017 | 0.040 |
| rs891511 | - | - | 0.086 | 0.025 | - | - | 0.103 | 0.029 | - | - | - | - |
| rs6969780 | - | - | 0.032 | 0.026 | - | - | - | - | - | - | - | - |
| rs76206723 | - | - | 0.035 | 0.026 | - | - | - | - | - | - | 0.036 | 0.026 |
| rs11977526 | - | - | -0.050 | 0.021 | - | - | - | - | - | - | -0.040 | 0.017 |

(continued from previous page)

| | SBP | | | | DBP | | | | PP | | | |
|---|---|---|---|---|---|---|---|---|---|---|---|---|
| rsid | Cluster | Mean | Estimate | SE | Cluster | Mean | Estimate | SE | Cluster | Mean | Estimate | SE |
| rs35783704 | - | - | 0.021 | 0.019 | - | - | 0.045 | 0.041 | - | - | 0.042 | 0.039 |
| rs2898290 | - | - | 0.001 | 0.016 | - | - | 0.002 | 0.040 | - | - | 0.002 | 0.027 |
| rs4454254 | - | - | 0.006 | 0.026 | - | - | - | - | - | - | 0.006 | 0.027 |
| rs34591516 | - | - | 0.033 | 0.024 | - | - | 0.065 | 0.048 | - | - | - | - |
| rs1449544 | - | - | 0.031 | 0.025 | - | - | - | - | - | - | 0.039 | 0.032 |
| rs72765298 | - | - | -0.020 | 0.026 | - | - | - | - | - | - | -0.024 | 0.030 |
| rs4245739 | - | - | - | - | - | - | 0.092 | 0.042 | - | - | - | - |
| rs1060105 | - | - | - | - | - | - | 0.024 | 0.043 | - | - | - | - |
| rs6429422 | - | - | - | - | - | - | -0.018 | 0.031 | - | - | - | - |
| rs7178615 | - | - | - | - | - | - | 0.042 | 0.044 | - | - | - | - |
| rs72799341 | - | - | - | - | - | - | -0.038 | 0.042 | - | - | - | - |
| rs1126464 | - | - | - | - | - | - | -0.037 | 0.031 | - | - | - | - |
| rs78378222 | - | - | - | - | - | - | -0.017 | 0.046 | - | - | 0.014 | 0.038 |
| rs745821 | - | - | - | - | - | - | 0.092 | 0.045 | - | - | - | - |
| rs6095241 | - | - | - | - | - | - | -0.082 | 0.045 | - | - | - | - |
| rs6108168 | - | - | - | - | - | - | 0.122 | 0.045 | - | - | - | - |
| rs9306160 | - | - | - | - | - | - | 0.151 | 0.034 | - | - | - | - |
| rs79146658 | - | - | - | - | - | - | 0.034 | 0.038 | - | - | -0.037 | 0.041 |
| rs4952611 | - | - | - | - | - | - | -0.016 | 0.044 | - | - | - | - |
| rs2579519 | - | - | - | - | - | - | -0.011 | 0.039 | - | - | 0.011 | 0.041 |
| rs2306374 | - | - | - | - | - | - | 0.349 | 0.042 | - | - | - | - |
| rs12374077 | - | - | - | - | - | - | -0.009 | 0.046 | - | - | - | - |
| rs9810888 | - | - | - | - | - | - | 0.038 | 0.045 | - | - | - | - |
| rs6825911 | - | - | - | - | - | - | 0.107 | 0.039 | - | - | - | - |
| rs66887589 | - | - | - | - | - | - | 0.208 | 0.045 | - | - | - | - |
| rs73030266 | - | - | - | - | - | - | 0.151 | 0.038 | - | - | - | - |
| rs10943605 | - | - | - | - | - | - | 0.113 | 0.036 | - | - | - | - |
| rs2071518 | - | - | - | - | - | - | -0.038 | 0.045 | - | - | 0.015 | 0.018 |
| rs76452347 | - | - | - | - | - | - | -0.017 | 0.046 | - | - | - | - |
| rs2289125 | - | - | - | - | - | - | - | - | - | - | -0.026 | 0.023 |
| rs2761436 | - | - | - | - | - | - | - | - | - | - | 0.047 | 0.034 |
| rs452036 | - | - | - | - | - | - | - | - | - | - | 0.000 | 0.027 |
| rs33063 | - | - | - | - | - | - | - | - | - | - | -0.038 | 0.034 |
| rs2645466 | - | - | - | - | - | - | - | - | - | - | 0.055 | 0.035 |
| rs740698 | - | - | - | - | - | - | - | - | - | - | 0.047 | 0.039 |
| rs28427409 | - | - | - | - | - | - | - | - | - | - | 0.058 | 0.034 |
| rs36010659 | - | - | - | - | - | - | - | - | - | - | 0.021 | 0.039 |
| rs9662255 | - | - | - | - | - | - | - | - | - | - | 0.041 | 0.038 |
| rs6081613 | - | - | - | - | - | - | - | - | - | - | -0.046 | 0.030 |
| rs7255 | - | - | - | - | - | - | - | - | - | - | -0.074 | 0.033 |
| rs9479200 | - | - | - | - | - | - | - | - | - | - | 0.014 | 0.031 |
| rs449789 | - | - | - | - | - | - | - | - | - | - | 0.180 | 0.036 |
| rs1322639 | - | - | - | - | - | - | - | - | - | - | 0.004 | 0.035 |
| rs1953126 | - | - | - | - | - | - | - | - | - | - | 0.059 | 0.037 |
| rs10818775 | - | - | - | - | - | - | - | - | - | - | 0.031 | 0.033 |

| | Effect | Genetic association with... | | | | | | |
|---|---|---|---|---|---|---|---|---|
| Genetic variant | allele | SBP | DBP | PP | CAD risk | Trunk fat % | Impedance of arm | Arm fat % |
| rs17249754 | G | 0.802 (0.062) | 0.408 (0.037) | 0.388 (0.042) | -0.055 (0.007) | -0.005 (0.003) | 0.002 (0.002) | -0.005 (0.002) |
| rs1438896 | T | 0.228 (0.050) | 0.191 (0.029) | 0.045 (0.034) | -0.020 (0.006) | 0.003 (0.002) | -0.004 (0.002) | 0.001 (0.002) |
| rs4494250 | A | 0.256 (0.049) | 0.177 (0.029) | 0.062 (0.033) | -0.015 (0.006) | 0.008 (0.002) | 0.010 (0.002) | 0.004 (0.002) |
| rs35410524 | T | 0.300 (0.059) | 0.141 (0.035) | 0.162 (0.040) | -0.032 (0.007) | 0.003 (0.003) | -0.004 (0.002) | 0.003 (0.002) |
| rs11030119 | G | 0.170 (0.051) | 0.133 (0.030) | 0.053 (0.034) | -0.016 (0.006) | -0.023 (0.002) | 0.013 (0.002) | -0.022 (0.002) |
| rs2304130 | G | 0.307 (0.086) | 0.234 (0.051) | 0.067 (0.058) | -0.028 (0.010) | -0.020 (0.004) | -0.007 (0.003) | -0.014 (0.003) |
| rs62104477 | T | 0.087 (0.049) | 0.130 (0.029) | -0.026 (0.033) | -0.016 (0.006) | 0.018 (0.002) | -0.007 (0.002) | 0.016 (0.002) |
| rs918466 | G | 0.139 (0.048) | 0.164 (0.028) | -0.042 (0.032) | -0.017 (0.005) | -0.010 (0.002) | -0.009 (0.002) | -0.006 (0.002) |
| rs687621 | A | 0.098 (0.048) | 0.133 (0.029) | -0.026 (0.033) | -0.027 (0.005) | -0.013 (0.002) | -0.015 (0.002) | -0.013 (0.002) |
| rs4387287 | A | 0.195 (0.062) | 0.167 (0.037) | 0.020 (0.042) | -0.033 (0.008) | - | - | - |

Table A2: Genetic associations (beta-coefficients and standard errors) for variants in the cluster with negative causal effect of blood pressure on coronary artery disease risk. Genetic associations with blood pressure (mmHg) were estimated in 299 024 participants of European ancestry from the International Consortium for Blood Pressure. Genetic associations with coronary artery disease (CAD) risk (log odds ratios) were estimated in 122 733 cases and 424 528 controls primarily of European descent from the CARDIoGRAMplusC4D consortium and UK Biobank. Genetic associations with the adiposity traits (SD units) were estimated in 337 199 participants of European descent from UK Biobank (Ben Neale estimates). Associations in UK Biobank for rs4387287 were not available.
